# Ribo-ITP enables identification of translons from limited input samples

**DOI:** 10.1101/2025.08.04.668486

**Authors:** Vighnesh Ghatpande, Uma Paul, Logan Persyn, Yifan Tian, MacKenzie A Howard, Can Cenik

## Abstract

In the last decade, an unexpectedly large number of translated regions (translons) have been discovered using ribosome profiling and proteomics. Translons can act as regulatory elements or encode functional micropeptides. However, identification of translons has been limited to cell lines or large organs due to high input requirements for conventional ribosome profiling and mass spectrometry. Here, we address this input limitation using Ribo-ITP on difficult-to-collect samples such as microdissected hippocampal tissues and single preimplantation embryos to identify thousands of translons. To test the translational capacity of the identified translons, we engineered a translon-dependent GFP reporter system and detected expression of translons initiating at ATG and near-cognate start codons in mouse embryonic stem cells (mESCs). Mutating the translons in mESCs identified a small proportion that may negatively impact growth. We identified distinct expression patterns of translons using a comparative analysis of more than a thousand ribosome profiling datasets across a wide range of cell types. Further, using a machine learning model we predict that specific upstream translons in synaptically enriched mRNAs regulate translation efficiency of the annotated coding region. Taken together, we present a proof-of-concept study to identify non-canonical translation events from low input samples which can be applied to cell and tissue types inaccessible to conventional methods.

## Introduction

Emerging evidence over the last decade strongly indicates pervasive translation in regions of the genome previously annotated as non-protein coding. To systematically investigate these non-canonical translation events, researchers have relied on ribosome profiling and its variants^1,2^ which remain the primary tools for discovery, complemented by mass spectrometry based approaches for peptide-oriented identification^3–5^. Nomenclature of such non-canonical translation events is often associated with the term ‘ORF’, which remains a point of confusion. In this study, we adopt the term ‘translon’ to refer to ORFs with evidence of translation^6^. While the annotated protein-coding region in a transcript is also a translon, we will refer to it as the CDS for clarity.

Translons and their encoded peptides can function in multiple cellular processes^7–21^, including signaling, chromatin remodeling, and metabolism. To determine the potential function of the encoded peptides, a variety of experimental methods have been developed, including CRISPR-Cas9 pooled knockout screens, reporter assays, phylogenetic analyses, and affinity-ligation based techniques^18,22–27^. Together, these complementary approaches have revealed that non-canonical translation occurs extensively and in a context-dependent manner across diverse cell lines and tissues.

Despite these advances, the catalog of micropeptides and translons from rare or difficult-to-isolate cell types remains incomplete^28^. Advances in low-input and single-cell technologies have uncovered important biological phenomena, including heterogeneity in cellular states^29,30^, tissue responses^31^, rare cell populations^32^, and developmental lineage tracing^33,34^. These approaches have been especially useful for identifying cell type specific gene expression patterns in complex diseases^31,35^. Yet, single-cell RNA-seq measures RNA abundance rather than translation, and conventional ribosome profiling and mass spectrometry require large amounts of input material. As a result, non-canonical translation in rare and difficult to isolate cell types remains almost entirely uncharacterized.

To address this gap, recent efforts have sought to improve ribosome profiling by reducing input requirements^36,37^. Among these, RIBOsome profiling by IsoTachoPhoresis (Ribo-ITP)^36^ is particularly promising. By combining isotachophoresis based RNA concentration with microfluidic size selection, Ribo-ITP minimizes sample loss and efficiently captures ribosome-protected fragments from as few as a single cell, providing high transcript coverage. This combination of sensitivity and scalability makes Ribo-ITP uniquely capable of uncovering translational events in rare or heterogeneous cell populations.

Activity-dependent translational changes in neurons are essential for synaptic plasticity^38–42^, yet ribosome profiling of LTP-induced acute hippocampal slices has been limited by the technical difficulty of scaling experiments to meet tissue input requirements^43^. Similarly, ribosome profiling in zebrafish from hundreds of embryos at different developmental stages enabled discovery of Toddler, a peptide essential for gastrulation and normal development^7^. However, similar applications in mammalian systems are challenging due to the cost and ethical concerns of collecting comparably large numbers of embryos, leaving non-canonical translation in these contexts uncharacterized.

Here, we present a proof-of-concept study applying Ribo-ITP to microdissected CA1 subregions of single acute hippocampal slices and 16- and 32-cell stage mouse preimplantation embryos. Previous Ribo-ITP studies uncovered translational regulation from single mouse embryos^36,44^, however, translon identification was hampered by nucleotide-level resolution loss in MNase-treated samples. Applying Ribo-ITP using RNase I, we detect thousands of translons expressed in these samples, demonstrating the potential of low-input ribosome profiling to map non-canonical translation in cell types and tissues previously inaccessible to conventional methods.

## Results

### Ribo-ITP enables identification of non-canonical translation events from microdissected mouse hippocampus

We applied Ribo-ITP to measure translation in microdissected mouse hippocampal CA1 regions. We induced LTP by theta burst stimulation and manually microdissected and flash-froze the potentiated region at 10 mins, 30 mins, and 60 mins post-LTP induction along with unstimulated control tissue (Fig1A, S1F). This stimulation activates two major types of neurons in this region, CA1 pyramidal neurons (∼89%) and GABAergic interneurons (∼11%). Our dissection methods further increase the proportion of pyramidal neurons but as some interneuron cell bodies lie in the stratum pyramidale layer, they can not be fully excluded^45^. Based on the dimensions of the microdissected region (see Methods), we estimate that these slices would contain a few thousand cells.

**Fig 1:**
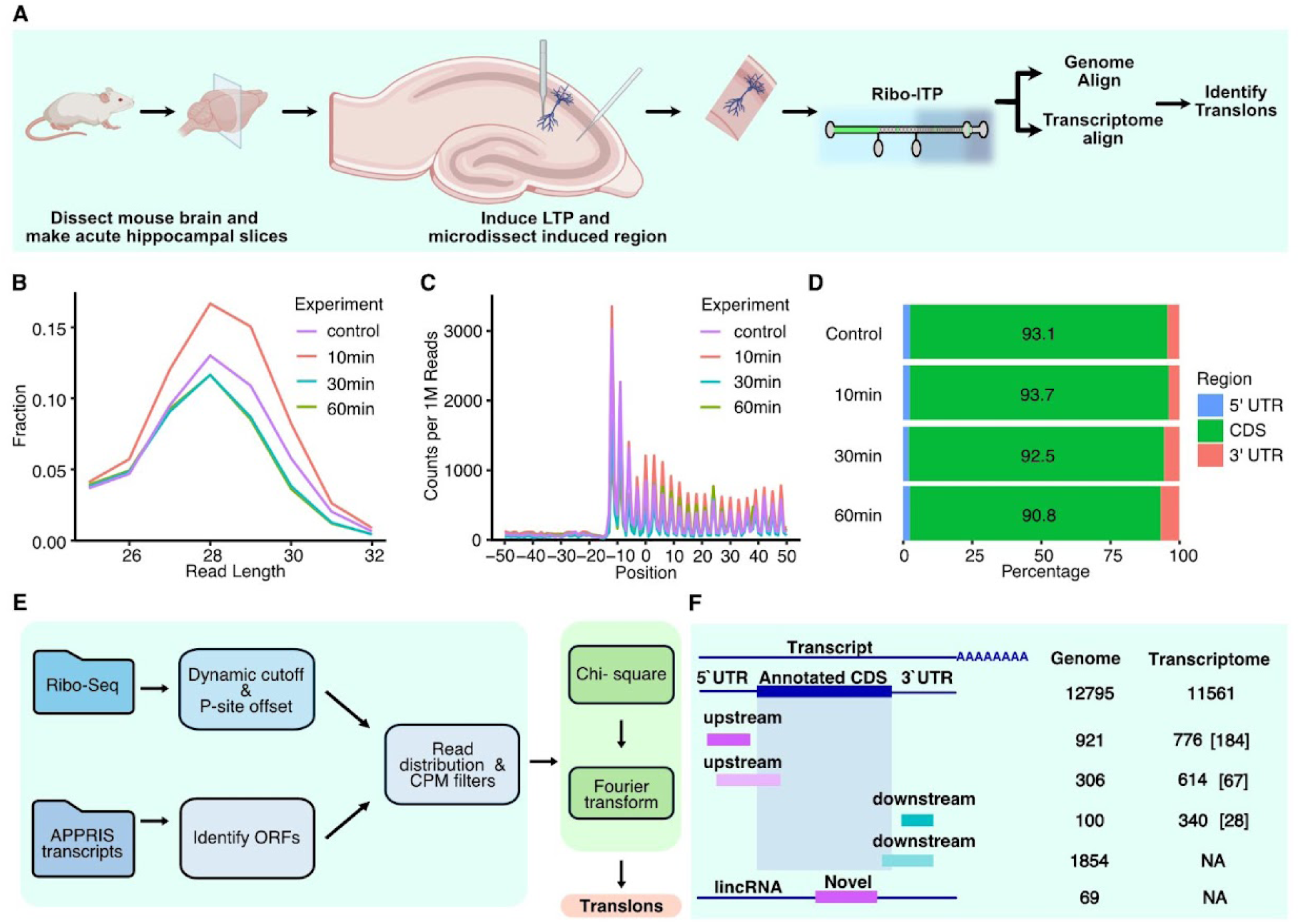
Ribo-ITP enables identification of non-canonical translation events from low input tissue samples A, Schematic of the experimental procedure for performing Ribo-ITP of LTP-induced microdissected CA1 region from acute slices of mouse hippocampal sections. B, Normalized length distribution of the transcriptome-mapping RPFs after merging replicates by time point. C, Ribosome occupancy around the translation start site after merging replicates by time points. Translation start site is denoted by the position 0. Aggregated read counts (y-axis) relative to the start site are plotted using the 5′end of the read. D, Distribution of reads across transcript regions (5′ UTR, CDS, and 3′ UTR) are shown. E, Schematic of the transcriptome-based translon identification strategy developed in this study. F, Schematic showing the different types of translons identified (left) and the corresponding number of ORFs identified for each event using the genome- and transcriptome-based approach. For transcriptome-based approach, the chi-square filtered counts are shown, with the number of Fourier transform (FT) filtered translons mentioned in brackets

Ribo-ITP resulted in libraries with ribosome protected fragment (RPF) length distributions centered at 28 nucleotides (nt) and were enriched at annotated start and stop codons (Fig1B, C, S1A-C). Most RPFs mapped to the annotated coding regions of transcripts (Fig1D, S1D) with >77% mapping to the dominant frame (FigS4D). The number of RPFs mapping to annotated coding regions across all 12 replicates was reproducible (Spearman correlation coefficient > 0.83; FigS1E). These results suggest that Ribo-ITP enables isolation of RPFs from microdissected, LTP-induced CA1 regions from single slices.

To determine the translational changes in the CA1 region, we analyzed expression patterns across time points. We did not observe consistent LTP dependent temporal changes of early immediate genes across replicates. While technically reproducible, the highly variable expression patterns across timepoints were likely attributable to variability in microdissection of the LTP-activated region. A more precise anatomical targeting and increased temporal and biological replicates would be required to identify statistically significant LTP-induced temporal changes.

Given the variability in temporal changes we performed a pooled analysis of all the timepoints. Pooling the deduplicated reads from 12 replicates resulted in over 20 million genome mapping and 22 million transcriptome mapping unique molecules which were used for identifying translons (Methods). Genome mapping is the most commonly used approach as it helps identify a broader range of translation events. However, RPFs are short (∼28nt) and can map to multiple regions in the genome, increasing mapping errors. Transcriptome mapping, on the other hand, can identify only a subset of translons but is less prone to mapping errors.

We first used RiboTISH, a genome-based pipeline and identified a total of 19,475 translons^46^. Out of these, 12,795 were known CDSs, and the remaining 6,680 were novel translons with respect to the annotations used. Of the novel translons, 1,227 initiated upstream, and 1,954 initiated downstream to the CDS (Fig1F), 69 were encoded in long intergenic non-coding RNAs (lincRNA), 3,265 were truncations of CDSs, 99 were internal but out of frame with respect to the CDS, 5 were from antisense transcripts, 5 were from putative protein coding transcripts, 3 were from small non-coding RNA transcripts, 51 were from pseudogenes, and 2 were from intronic regions of genes (TableS1).

We next developed a transcriptome-based approach which uses a .ribo file^47^ as the input for translon detection (Fig1E; Methods and see FigS4A,C for details of filtered-out translon counts and the start codon distribution at each step). We first assessed the periodicity of RPFs on a given translons using a chi-square test (Methods) and identified 1,390 translons upstream and 340 downstream of annotated CDS, along with 12,795 expressed CDSs (more than 1 CPM reads). We then assessed the expression and periodicity of footprints using a database (RiboBase) of over ∼2.35 billion transcriptome mapping reads from ∼1,600 highly periodic mouse ribosome profiling datasets from 41 different cell and tissue types (see Methods)^48^. Given the high coverage of transcripts within these datasets, we applied a Fourier transform to further assess the periodicity of RPFs mapping to the translons that passed the chi-square test (see Methods). 251 upstream ORFs and 28 downstream ORFs had evidence of periodic RPF reads in the database (Fig1F, TableS2). 34% of transcriptomic translons overlapped with the genomic translons with no clear bias towards the amino acid length but ATG initiating and upstream non-overlapping translons were overrepresented (FigS4E-G). Furthermore, translons on annotated long non-coding RNAs could not be detected as our reference transcriptome only contained protein coding transcript isoforms. These results highlight the significant impact of the choice of computational pipeline on detection of translons in agreement with recent studies that reported poor overlap between the set of translon calls from different computational methods^49^.

To understand the impact of the various filtering steps in our transcriptomic pipeline, we assessed the percentage of translons that were filtered at each step. We found that the Fourier transform filter was the most stringent, filtering out the largest number of transcripts in the hippocampal dataset, while none of the filters introduced a bias towards ATG start codon (FigS4A,C). Taken together, these results establish that thousands of non-canonical translons can be detected from microdissected CA1 regions from single hippocampal acute slices using Ribo-ITP.

### Ribo-ITP enables identification of non-canonical translation events from single mouse embryos

We tested if our approach would help identify translons from even fewer cells and employed Ribo-ITP to profile single 16- and 32-cell stage preimplantation mouse embryos (Fig2A). We treated stage-matched embryos with Harringtonine, a translation inhibitor which stalls initiating and early elongating ribosomes^50,51^. In total, we profiled 15 single embryos from 16-cell and 32-cell stages using Ribo-ITP. These libraries had RPF length distributions centered around 28-30 nt (Fig2A-D) and showed enrichment at annotated start codons (S2E-H). Most of the RPFs mapped to the annotated coding regions of transcripts (Fig2B), with 62% RPFs mapping to the dominant frame (FigS4D). Upon Harringtonine treatment, reads mapping in the 5′ UTR junction, i.e. 50 nt surrounding the annotated CDS start site, increased at least 2-fold (Fig2C, S2J,K).

**Fig 2:**
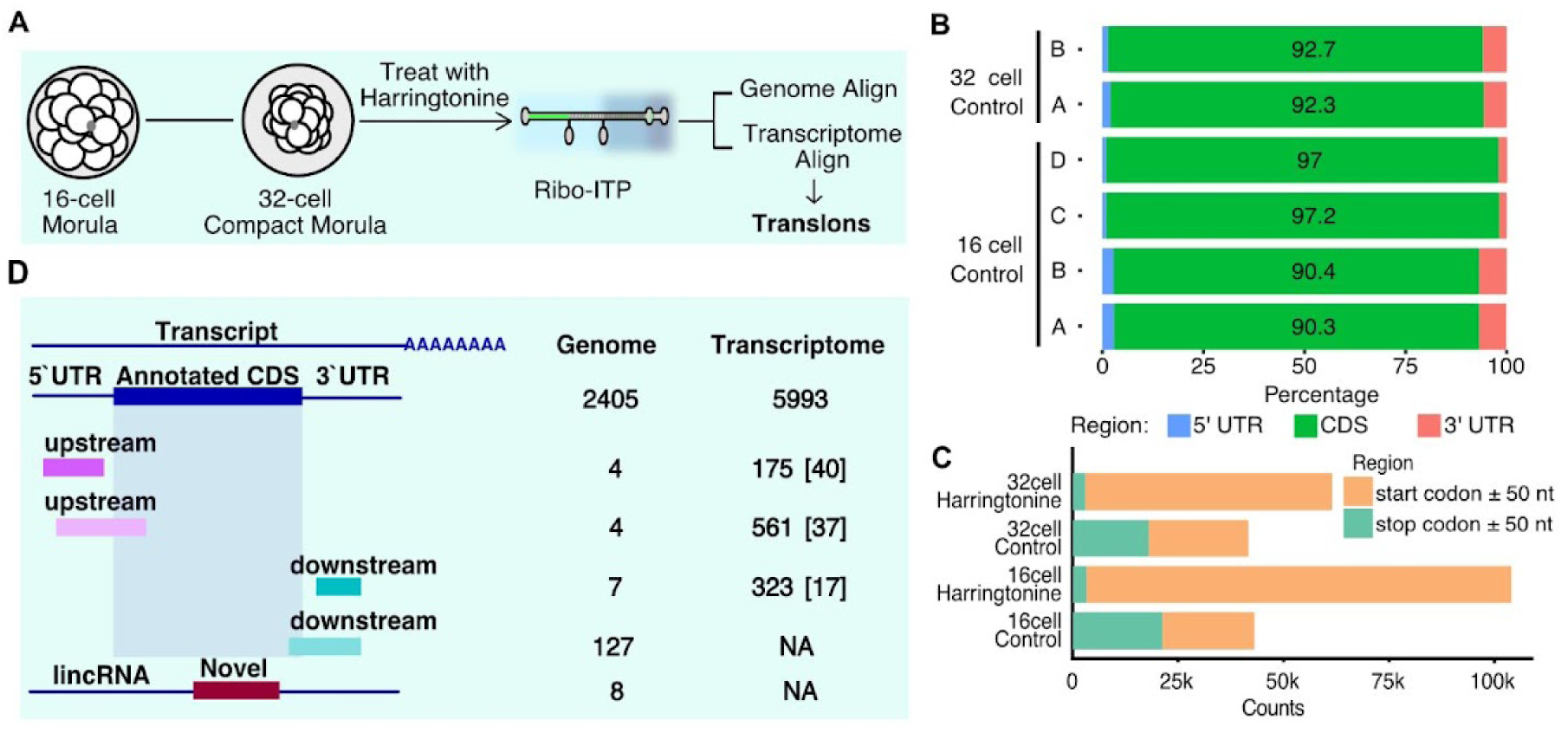
Ribo ITP enables identification of non-canonical translation events from single mouse embryo A, Schematic of the experimental procedure for performing Ribo-ITP of 16- and 32-cell single embryos. B, Percentage of reads mapping to the different transcriptomic regions (5′UTR, CDS, and 3′UTR) in 16-cell and 32-cell control embryos. C, Distribution of reads mapping in the 50 nt region surrounding the translation start site. The raw counts are merged from multiple replicates per stage. D, Schematic showing the different types of events identified (left) and a table of corresponding number of ORFs identified for each event using the genome- and transcriptome-based approach. Number of FT filter pass translons are mentioned in brackets.

Pooling all the untreated libraries together, we mapped over half a million unique RPFs to both the genome and transcriptome. We used RiboTISH and identified 2,555 translons, of which 2,405 were CDSs, 8 initiated upstream to the CDS, 134 initiated downstream to the CDS, and 8 were encoded in lincRNAs (Fig2D, TableS3). Our transcriptome-based approach identified 7,052 translons out of which 5,993 were CDSs (>1 CPM), 736 translons upstream and 323 downstream of annotated CDSs (TableS4). These translons passed the chi-square 3-nt periodicity filter (Fig1E, FigS3B,C). Out of these, 66.7% upstream translons and 28.5% downstream translons had RPFs mapping around the predicted start site (±2 codons) in the Harringtonine-treated datasets. Fourier transform based analysis identified high periodicity across 77 upstream translons and 17 downstream translons with consistent periodicity across studies in RiboBase (Fig2D). Nonetheless, candidates that do not pass the Fourier transform filter may also represent bona fide translons, but validating their expression would require further experimentation. Taken together, these findings suggest that Ribo-ITP can be used to identify translons from single preimplantation mouse embryos.

### Properties of identified translons

To characterize the physicochemical properties of the identified translons, we first analyzed their length distributions. In hippocampal samples, transcriptome-annotated upstream and downstream translons exhibited median lengths of 46 and 51 amino acids (aa), respectively (Fig3A). Genome-annotated translons showed greater variability, with median lengths of 53 aa for upstream translons, 150 aa for downstream translons, and 65 aa for lincRNA translons (FigS4A). Due to the limited number of genome-annotated translons in embryonic samples, subsequent analyses focused on transcriptome-annotated translons, which displayed median lengths of 35 aa and 31 aa for upstream and downstream translons, respectively (Fig3D).

**Fig 3:**
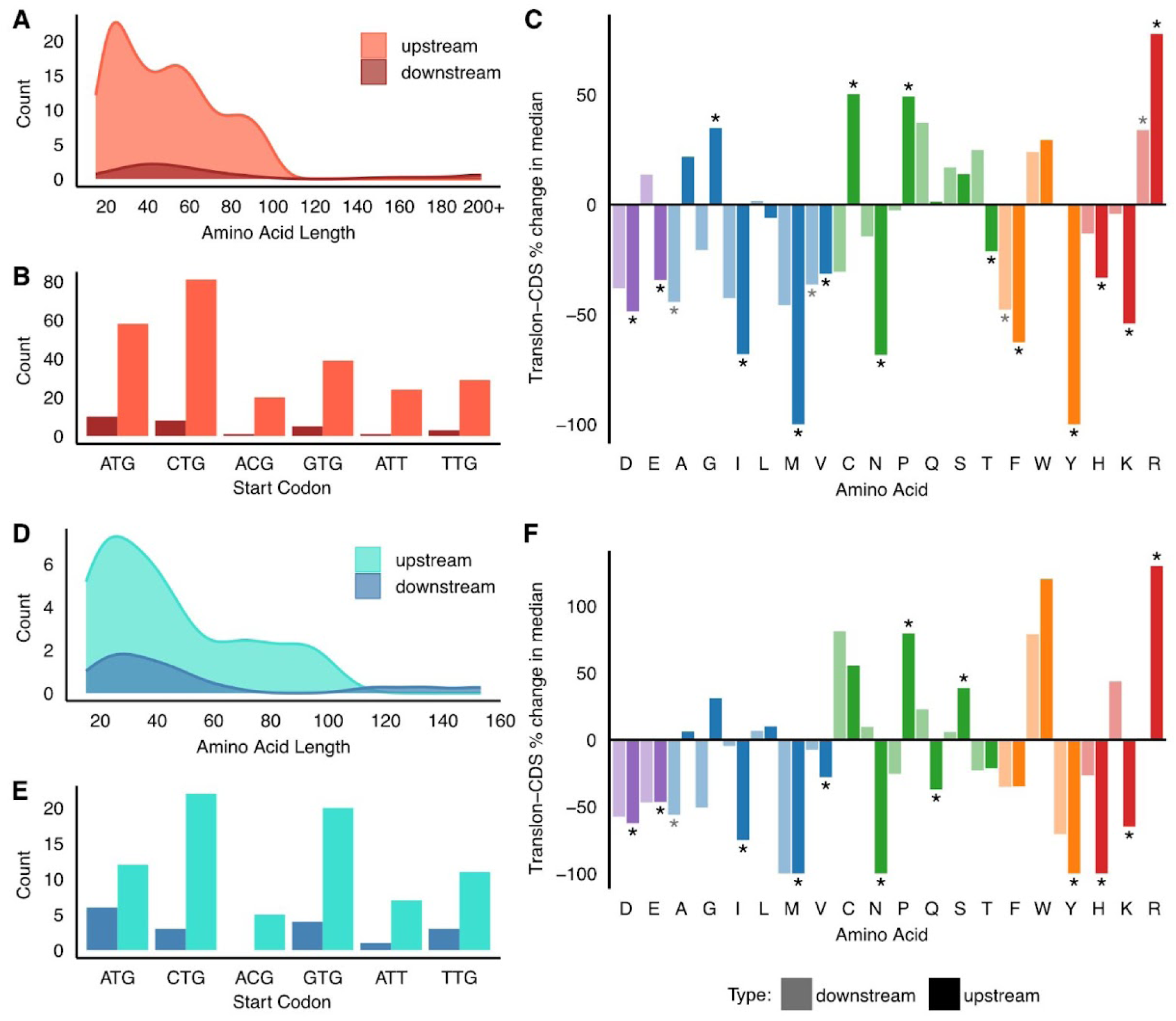
Properties of embryonic and hippocampal translons. A, Density plots showing the amino acid length distribution of the peptides encoded by the translons identified in the hippocampal dataset. B, The number of translons identified with cognate and near-cognate start codons in the hippocampal dataset. C, Bar graph showing the percent change in the median amino acid frequency of each residue when comparing upstream (dark) or downstream (light) hippocampal translons to the CDS. Colors indicate amino acid type. * indicates Wilcoxon’s rank-sum test p-values < 0.05. D, Density plots showing the amino acid length distribution of the peptides encoded by the translons identified in the embryo dataset. E, The number of translons identified with cognate and near-cognate start codons in the embryo dataset. F, Bar graph showing the percent change in the median amino acid frequency of each residue when comparing embryonic upstream (dark) or downstream (light) translons to the CDS. Colors indicate amino acid type. * indicates Wilcoxon’s rank-sum test p-values < 0.05.

Analysis of initiation codon revealed distinct patterns between translon categories. In both hippocampal and embryonic datasets, the majority of transcriptome-annotated upstream translons initiated at CTG, a near-cognate start codon, whereas downstream translons predominantly initiated at the canonical ATG codon (Fig3B,E). Genome-annotated translons in hippocampal samples preferentially used ATG as their initiation codon (FigS4B).

Given that the amino acid composition influences peptide stability^52^, we compared the composition of the translon-encoded peptides with the CDS encoded protein in the same transcript (Methods). Although, the amino acid composition of the translon-encoded peptides was similar between both the neuronal (see Fig3C for transcriptome-annotated, S4C for genome-annotated) and embryo translons (Fig3F), they had a significant bias for specific amino acids as compared to the CDS. Particularly, in upstream translons, the amino acids arginine and proline were significantly enriched (*p-value*: hippocampal (h) _Arg_ = 2.20×10^-21^, embryonic (e) _Arg_=3.93×10^-9^, h_Pro_=0.00326, e_Pro_=0.00621) while aspartic acid, glutamic acid, asparagine, isoleucine, methionine, valine, tyrosine, histidine, and lysine were significantly depleted (*p-value*: h_Asp_=1.30×10^-16^, e_Asp_=8.22×10^-9^, h_Glu_=1.77×10^-13^, e_Glu_=7.13×10^-7^, h_Asn_=4.61×10^-24^, e_Asn_=6.20×10^-10^, h_Ile_=3.26×10^-24^, e_Ile_=5.77×10^-6^, h_Met_=4.88×10^-8^, e_Met_=0.00144, h_Val_=2.91×10^-14^, e_Val_=0.0400, h_Tyr_=2.12×10^-32^, e_Tyr_=1.52×10^-10^, h_His_=0.00278, e_His_=0.0288, h_Lys_=9.84×10^-17^, e_Lys_=1.28×10^-10^). These findings align with observations from post-mortem human brains,^15^ where the translons were enriched in aromatic amino acids and arginine but depleted in polar amino acids and lysine. Downstream translons exhibited significant alanine depletion in both datasets (*p-value*: h= 0.0162, e=0.0284). Taken together, these results suggest that translon-encoded peptides possess distinct physicochemical properties compared to annotated CDS products.

### Non-canonical translons can drive translation of a reporter and affect cellular proliferation

To assess the functional capacity of identified translons, we initially tested whether these sequences possess translational capacity to drive the expression of a reporter construct. We selected seven embryonic upstream translons and cloned each sequence from the transcript start to the stop codon of the translon in frame with GFP lacking its start codon (FigS5A). This design ensures that fluorescence is detectable only when the translon is robustly translated.

We included three controls: (1) GFP with its native ATG start codon (start-GFP) as a positive control, (2) GFP lacking a start codon (nostart-GFP) as a negative control, and (3) insulin transcript (5′ UTR + CDS), which codes for a 110 amino acid pro-peptide in frame with nostart-GFP. Upon expression in mESCs, the positive control had robust GFP expression (FigS5B), the negative control showed no detectable fluorescence (FigS5C), and the insulin control displayed weak GFP expression (FigS5D), likely due to pro-peptide cleavage and the tissue-specific nature of insulin expression.

Four of the seven tested embryonic translons drove variable levels of GFP expression. Due to the technical challenges of quantifying fluorescence intensity of the three-dimentional mESC colonies with variable cell numbers, we decided to present these results qualitatively. Translons within *Eif5* and *Clpx* mRNAs produced strong GFP expression (FigS5E,F), while translons on *Npm1* and *Hdgf* mRNAs drove weak fluorescence (FigS5G-H). Translons on *Rps12*, *Olfr1217* and *Top1* mRNAs did not show any GFP expression (FigS5I,K). Notably, the Eif5 translon represents the most highly expressed upstream translon in 16-cell and 32-cell embryos and is a well-characterized regulatory translon^46,53^. To the best of our knowledge, the upstream translon in the *Clpx* transcript has not been previously reported. This translon initiates at a CTG codon with no downstream in-frame ATG, indicating that translons with non-canonical initiation codons can drive robust translation.

Given the success of the GFP reporter strategy, we expanded our investigation to assess the translational potential of all the identified translons using a FACS-based pooled GFP reporter screen (Fig4A). We cloned 443 translons into the nostart-GFP lentiviral vector and performed a pooled infection of mESCs at a low multiplicity of infection. GFP+ cells (∼5% of total cells, FigS5L) were sorted and the integrated translons were identified by sequencing. We detected 378 of 443 (85%) tested translons in the GFP+ population, with high reproducibility across replicates (Spearman’s ρ = 0.85, FigS5Q). This result suggests that a substantial portion of the identified translons can drive GFP reporter translation.

**Fig 4:**
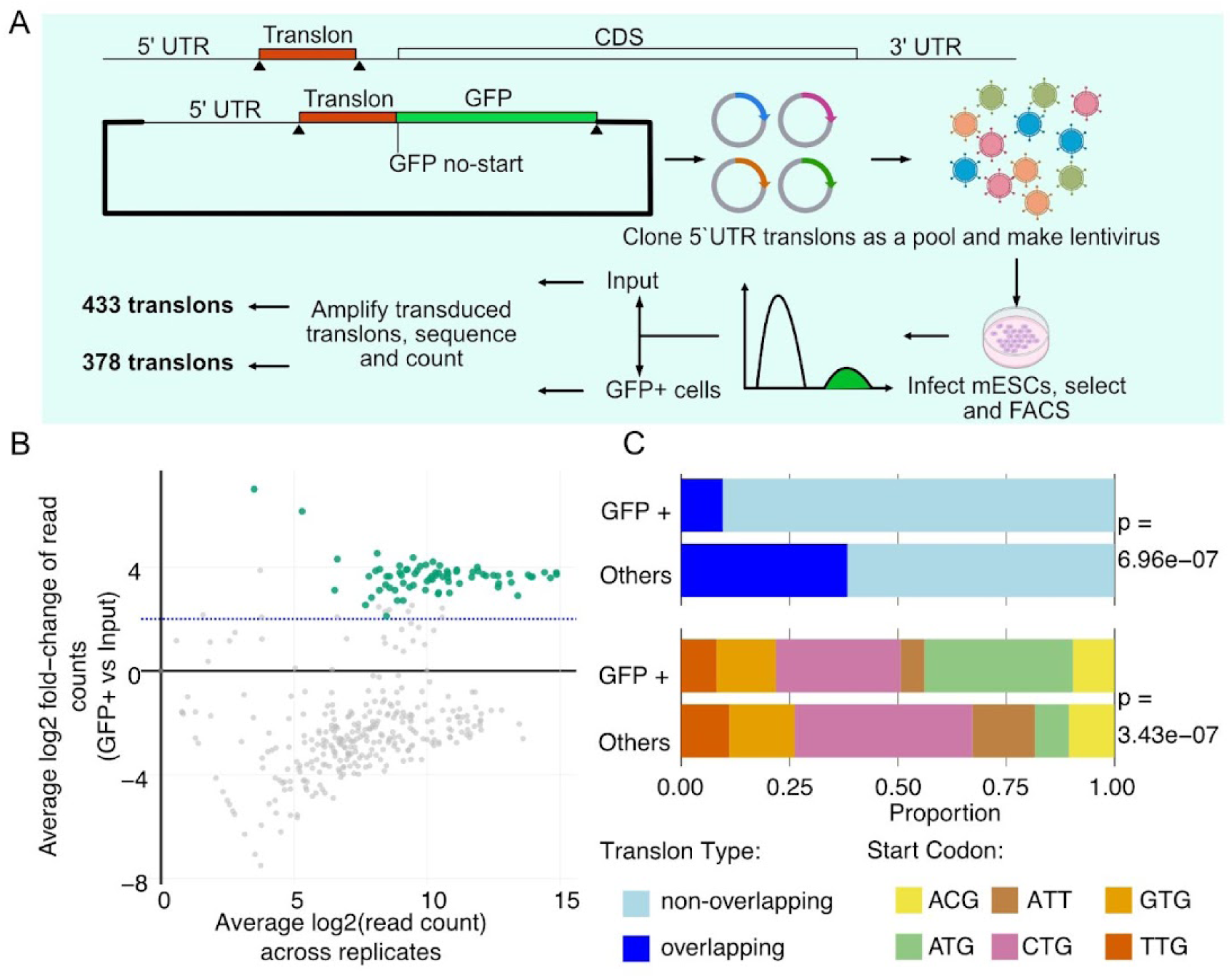
Translons drive translation of GFP reporter. A, Schematic of the reporter design for GFP-based translational capacity reporter. B, MA plot showing the fold change of read counts in GFP+ cells versus input cells and average read counts across replicates. Dotted blue line indicates average log_2_FC = 2. The translons showing 4-fold enrichment in both the replicates are colored in green while the remaining translons are colored grey. C, Stacked bargraphs showing proportion of translons annotated by either translon type or start codon between the GFP+ and other populations. For significance testing, Wilcoxon’s sum rank test was used for translon type and Fisher’s exact test p-value for start codon distribution.

We reasoned that the low percentage of GFP+ cells (FigS5L) likely indicates that most translons likely exhibit heterogenous expression across cells. To test this hypothesis, we transfected mESCs with equal amounts of either the startGFP plasmid or 4 randomly selected library clones. As expected, startGFP plasmid drove robust GFP expression with high transfection efficiency (Fig S5M, GFP+ vs phase contrast). In contrast, the clones showed heterogeneous GFP expression (FigS5N-P, GFP+ vs phase contrast).

To identify translons with the strongest translational activity, we compared the mean log_2_ fold change (log_2_FC) between input and GFP+ population and defined 73 translons as “GFP-enriched” based on exhibiting both mean log_2_FC and mean log_2_ expression > 2 in both the replicates (Fig4B). Comparison of GFP-enriched translons to all others revealed distinct sequence features. In the GFP-enriched population, ATG initiation codon was significantly overrepresented (34% vs 7%, p= 3.43X10^-7^) (Fig4C). Additionally, translons that did not overlap annotated CDSs were significantly enriched among GFP-enriched translons (90% vs 61%, p= 6.96X10^-7^). Together, these results reveal that while most tested translons (378/443, 85%) show detectable translation activity, robust translation is driven by only a small subset (73/443, 16%), characterized by ATG initiation codons and lack of overlap with the CDS.

Out of the 73 GFP-enriched translons, 63 were detected in hippocampal datasets, 8 in embryonic datasets and 1 in both. The GFP-enriched embryonic translons were harbored in transcripts spanning diverse biological processes: *Rps12* is a ribosomal protein, *Huwe1*^54^ is involved in proteasomal degradation, Spns1^55^ localizes to lysosomes, *Ap2m1*^56^ functions in vesicle trafficking, and *Zbtb17*^57^ is chromatin regulator. In contrast, many of the translon-containing transcripts identified in the hippocampal data were involved in brain and neuronal development as well as synaptic signalling. Specifically, *Cacng8* and *Neto1* directly control kinetics of glutamate receptors expressed in the hippocampus^58,59^, while mutations in *Clcn4*, *Chrm4*^60^, *Gmp6a*^61^, and *Ncdn*^62^ are associated with epilepsy and intellectual disorders. These results suggest that translons may play regulatory roles in transcripts with established neuronal functions, making their functional characterization a compelling future direction.

Translons embedded in transcripts with essential cellular functions could themselves influence cellular fitness if they play a regulatory role. We therefore reasoned that mutating such translons in mESCs might result in measurable growth defects. We designed a pooled CRISPR-Cas9 screening strategy to introduce indels in 67 translons (see Methods). We aimed to introduce mutations targeting the region of the translons that is non-overlapping with the CDS. To detect subtle growth defects, mESCs were grown for approximately 20 doublings (day 10) after selecting for cells expressing gRNA and Cas9 (FigS5R). Perturbations in 4 out of the 66 translons led to subtle reductions in growth (*p*<0.05, FDR<0.25; FigS5S, TableS7). Although only a small fraction of tested translons affected growth, these findings suggest that a subset of translons may contribute to cellular fitness.

### Expression patterns of non-canonical translons

Beyond individual functional validation, expression patterns across tissues and cell types can reveal translons with potential regulatory roles or context-dependent functions. We therefore used RiboBase^48^, a compendium of ribosome profiling datasets spanning diverse cell types and tissues, to assess the expression of the identified translons. Embryonic upstream translons had robust expression across most tissues and cell types examined (>10 CPM median RPFs in most cell lines; Fig 5A, Table S5). In contrast, embryonic downstream translons exhibited consistently low expression across samples (<1 CPM median RPFs in most cell lines; Fig 5A, Table S5). These patterns suggest that upstream translons identified in preimplantation embryos represent constitutively active translational events rather than developmentally restricted phenomena.

**Fig 5:**
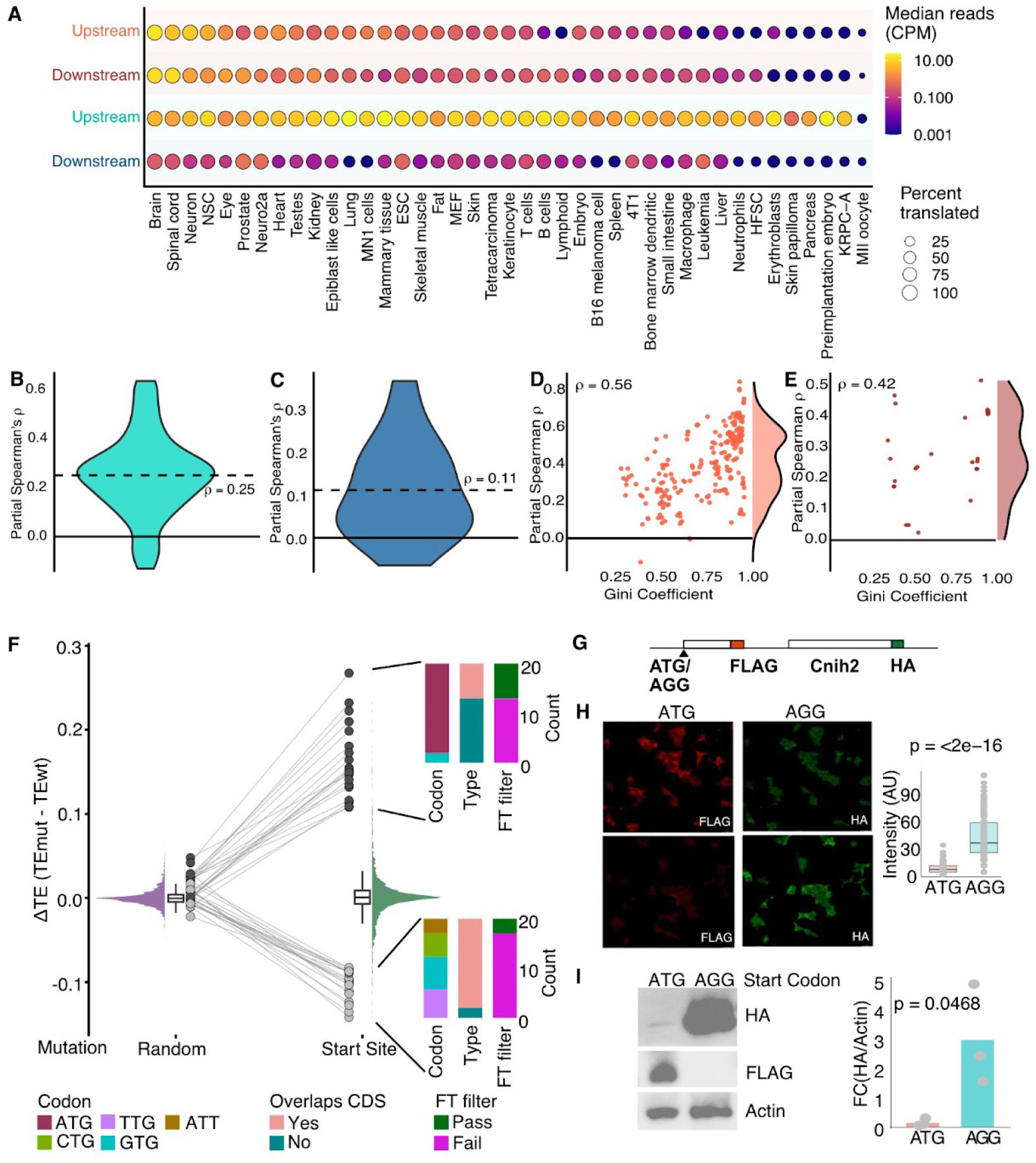
Expression patterns of translons A, The average expression of all the identified translons across the different cell types in RiboBase is plotted. The color coding of the dots represents the median expression of all the translons aggregated for each cell type and the size of the dot represents the percentage of expressed translons. The cooler color scheme on the y-axis represents the hippocampal translons while the warmer color scheme represents embryonic translons. B, Spearman’s ρ values for the partial correlation between the RNA normalized embryonic upstream translons and their corresponding CDS ribosome occupancy across all RiboBase studies. C, Spearman’s ρ values for the partial correlation between the RNA normalized embryonic downstream translons and their corresponding CDS ribosome occupancy across all RiboBase studies. D, Spearman correlation between the Spearman’s ρ values for the partial correlation between the RNA-normalized hippocampal upstream translons and CDS ribosome occupancy across all RiboBase studies with Gini coefficient. Distribution of Spearman’s ρ value for partial coefficients is shown on the right. E, Correlation between the Spearman’s ρ values for the partial correlation between the RNA normalized hippocampal downstream translons and CDS ribosome occupancy across all RiboBase studies with Gini coefficient. Distribution of Spearman’s ρ value for partial coefficients is shown on the right. F, Predicted change in TE_CDS_ upon mutating the start site of the translon versus a random position in the 5′UTR outside of the translon to a stop codon is plotted. The histograms visualize the distribution and the box plots denote the median and 90th percentile of the change in TE_CDS_. The top and bottom 20 translons are shown as dots with lines connecting them between the two conditions. The start codon, type and number of translons passing the FT filter are shown as stacked bar graphs. G, Schematic of the construct used for testing the effect of mutating the translon start codon on the CDS. H, Immunofluorescence images of HEK293T cells expressing the constructs with or without translon ATG stained with anti-Flag or anti-HA antibodies to detect expression of the translon and Cnih2. Intensity for HA signal in individual cells is quantified in the graph on the right. n=119 for AGG and n=115 for ATG-expressing cells. p-value indicates t-test results. I, Western blots of HEK293T cells expressing the constructs with or without translon ATG probed with anti-Flag or anti-HA antibodies to detect expression of the translon and Cnih2. Ratio of densitometric quantification for signal intensity of HA/Actin is plotted in the graph on the right. Ratios of 3 independent replicates are shown as grey dots. p-value is for unpaired t-test.

In contrast, hippocampal translons displayed more cell-type specific expression patterns. Both upstream and downstream translons had higher expression in primarily neuronal tissues (>10 CPM median RPFs in brain, spinal cord, neurons, neural stem cells), and in heart tissue (Fig5A, TableS5). The enrichment in neuronal contexts suggests potential tissue-specific roles for hippocampal translons compared to embryonic ones, though this difference may also be attributable to ∼40 times more RPFs in the hippocampal dataset compared to single embryos.

Upstream and downstream translons can modulate translation of their associated CDS^19,20,24,63^. To assess this relationship, we calculated the partial correlation between the ribosome occupancy of the CDS and the translon while accounting for RNA abundance. For embryonic upstream translons, we observed a modest positive partial correlation between the ribosome occupancy on the translon and the CDS (mean Spearman’s ρ = 0.25, Wilcoxon *p-value* = 1.81×10^-13^; Fig5B), consistent with previous observations in postmortem human brains and zebrafish embryos^15,19^. Embryonic downstream translons showed a weaker correlation with CDS occupancy (mean Spearman’s ρ= 0.11, Wilcoxon *p-value* = 7.63×10^-4^; Fig5C). Earlier reporter assays^63^ have demonstrated that the presence of a translated downstream open reading frame (dORF) enhances CDS translation efficiency. Specifically, they tested reporters containing zero, one, two, or multiple dORFs and found that CDS translation increased proportionally with dORF number when the dORFs were translationally active, but this enhancement was abolished when dORF translation was prevented by mutating their start codons^63^. However, the low ribosome occupancy of most downstream translons we identified in this study may limit our ability to detect such regulatory relationships.

Hippocampal translons exhibited stronger correlations with CDS expression than embryonic translons. Hippocampal upstream translons showed a mean partial correlation of ρ = 0.56 (Fig 5D-E; Wilcoxon *p* = 1.69×10^-41^), while downstream translons showed ρ = 0.42 (Wilcoxon *p* = 1.49×10^-8^). Notably, these correlations showed a bimodal distribution. We hypothesize that this may relate to the tissue-specific expression of these translons. Indeed, upstream translons with higher tissue specificity (higher Gini coefficients) exhibited stronger positive correlation with CDS translation (ρ vs. Gini, *p-value* < 2.2×10⁻¹⁶). For the downstream translons, this association was weaker but still significant (*p-value* = 0.02). Taken together, these results suggest that translational coordination between the upstream and downstream translons with the CDS is influenced by their expression specificity, with the hippocampal translons having functions in the nervous system.

Among the top 20 genes with high partial Spearman’s correlation coefficient were ionotropic receptor genes *Grin2b*, *Gria2* and *Gabra2* involved in neurotransmitter gated ion channel activity, as well as signalling intermediates like *Camkv*^64,65^ and *Mapk10*^66^, all of which are involved in synaptic transmission and signalling. In contrast, the top 20 genes with low partial Spearman’s correlation coefficient included *Sptbn2*^66,67^ and *Vps13d*^68^; mutations in which result in neurodegenerative disorders like spinocerebellar ataxia and amylotropic lateral sclerosis/frontotemporal dementia. Together, these results suggest that upstream translons in genes involved in neuronal processes and neurological diseases might be regulating CDS expression in an opposing manner.

Peptidomic evidence can provide further support for stability of the translon-encoded peptides. To identify if there are any embryonic translons with evidence of translation, we mined a proteomic dataset^69^ but we did not identify any peptides mapping to the translons we identified. Due to the often low abundance of micropeptides, their detection requires specialized protocols that enrich small proteins and peptides from cellular lysates^70,71^. Such a proteomic pipeline was not used in this study; thus, it is possible that micropeptides would have been missed. To identify if the hippocampal translons have peptidomic evidence, we mined two proteomic datasets focusing on non-canonical translation in mice brains and identified peptide evidence for a total of 47 translons. This included 45 translons from Wang et al.^72^, who profiled various embryonic and adult mouse organs, and 2 translons in Budamgunta et al.^73^, who profiled adult mouse striatum. These translons (TableS6) with proteomic evidence from independent studies are high confidence translons which might have functions in synaptic processes. Synaptic signalling is fundamentally driven by kinase- and phosphatase-dependent phosphorylation cycles, setting the stage for rapid activity dependent modulations^74–76^. Micropeptides have been previously identified to help in protein phosphorylation^77–80^. We identified multiple micropeptides originating from translons on mRNAs encoding kinases like *Map3k9*, *Map3k19*, *Wnk1,* and *Bmpr2* (21 mRNAs) and phosphatases like *Pten*, *Ppm1b*, *Dusp3*, and *Ppp1r9b* (9 mRNAs) to name a few; these could be tested for their role in regulating activity dependent signalling in synapses using targeted experiments.

### Subset of upstream translons regulate translation of the CDS

A scalable approach to assess whether a translon is regulatory is to predict how perturbing its translation affects CDS translation efficiency, rather than relying on low-throughput reporter assays^63,81,82^. Recent sequence-based models, including a multitask deep convolutional neural network (RiboNN), enable rapid prediction of CDS TE from transcript sequence alone^83,84^.

For each translon, we mutated *in silico* its start codon to a stop codon and, as a control, introduced the same stop codon at three random positions in the UTR outside the translon. The effect of each mutation was quantified as ΔTE (TE_Mut − TE_WT), where positive and negative values indicate increased or decreased CDS TE, respectively. Because some well-characterized regulatory transcripts, such as Gcn4^85^ and Atf4^82^ are translationally controlled only under stress conditions, we included all translons passing the chi-square filter rather than restricting the analysis to fourier transform passing translons.

Consistent with their role in modulating the ribosome loading^86–88^ on to CDSs, Disrupting upstream translons altered CDS TE (*F*(1972,1972) = 8.84, *p* < 0.001). In contrast, mutations in downstream translons had no detectable effect (Fig. S6A,B), likely reflecting the limited contribution of 3′UTR sequence on steady state TE^83^. To identify CDSs most sensitive to upstream translon perturbation, we analyzed the top and bottom 1% of ΔTE values (Fig. 5F). Most CDSs with increased TE harbored an ATG-initiating upstream translon (18/20), whereas all CDSs with decreased TE were associated with near-cognate-initiating upstream translons. Positive ΔTE translons were predominantly non-overlapping with the CDS (13/20), while negative ΔTE translons largely overlapped the CDS (Fig. 5F). Together, these results suggest that ATG-initiating, non-overlapping upstream translons tend to repress CDS translation, whereas overlapping translons initiating at near-cognate start codons may enhance CDS TE, potentially by slowing ribosome scanning and improving recognition of the CDS start codon.

To experimentally test these computational predictions, we selected the upstream translon in the *Cnih2* gene, which showed a strong predicted increase in CDS translation upon translon start-codon mutation (ΔTE = 0.198) compared to a minimal effect in control mutations (ΔTE = 0.023). *Cnih2* encodes an auxiliary subunit of AMPA-type ionotropic glutamate receptors that is highly expressed in the CA1 region of the hippocampus and regulates receptor trafficking and gating properties^89,90^. In an exogenous reporter construct, we tagged the upstream translon with FLAG and the CDS with HA, and mutated the translon start codon from ATG to AGG (Fig. 5G). Upon transfection into HEK293T cells, both western blotting and immunofluorescence analyses showed that disruption of translon initiation increased the protein expression of the *Cnih2* CDS (Fig. 5H,I), providing experimental validation of one of the computational predictions. Together, these results suggest that upstream translons identified from limited-input hippocampal and embryonic samples can function as regulatory elements controlling CDS translation. Such regulation may be particularly important in activity-dependent synaptic contexts, where rapid and precisely tuned translational responses are required.

## Discussion

We present proof-of-concept experiments demonstrating that Ribo-ITP enables the detection of translons in samples that are inaccessible to conventional ribosome profiling due to challenges in collecting sufficient quantities of material. This method yields high-quality, reproducible ribosome occupancy profiles with strong 3-nt periodicity, essential for identifying translons. Despite conventional ribosome profiling using orders-of-magnitude more input, of the 1,625 previously published ribosome profiling studies, only 20% achieve a similar or better 3-nt periodicity as we have in our datasets (FigS3D). While recent efforts proposed to build atlases of micropeptides and translons using conventional approaches^28^, such strategies will remain limited to abundant cell types and large tissues. The diversity of translons in rare, biologically important cell populations would therefore remain inaccessible. Our study provides a path forward for capturing non-canonical translation in these contexts.

In this study, we used Ribo-ITP to measure translation from the microdissected CA1 region of a single hippocampal slice, which is an orders-of-magnitude reduction in input compared to previous ribosome profiling studies that pooled ∼40-50 slices from multiple animals^38,39,43,91^. This reduced input lowers both experimental burden and cost. Similarly, previous studies in zebrafish required 400-600 embryos per replicate to identify micropeptides ^7,92^. Such large-scale collection is either impractical or currently impossible in many systems, including *C. elegans*^93^ and mammals^36^.

Our transcriptome-based translon detection pipeline allowed us to leverage a large compendium of uniformly processed ribosome profiling data (RiboBase)^48^. This resource contains paired data of ∼1600 mouse experiments from different cell and tissue types, enabling expression analysis of translons across diverse contexts. Most translons detected in embryos exhibited broad expression, while those identified in the CA1 region of the hippocampus were largely specific to the nervous system. This analysis suggests that translons may support both housekeeping functions and cell type-specific roles. These observations raise questions about the regulation and function of tissue-specific translons: What mechanisms underlie their specificity? Do they contribute to cellular homeostasis? Expanding the compendium of ribosome profiling datasets will likely enable uncovering the full diversity and biological significance of these translons.

The peptides that may be encoded by the identified translons were enriched for proline and arginine and depleted in methionine, aspartic acid, histidine, and lysine. Noncanonical proteins and canonical protein isoforms starting with non-methionine amino acids are enriched for degradation-related features like high C-term hydrophobicity and C-term degrons compared to all canonical proteins^94^. Arginine in the C-terminal can act as a degron leading to degradation of the protein^95^. Stop codon read-through leads to translation into the 3′UTR, resulting in proteins with 3′ UTR-encoded peptides or poly-lysine tracks that destabilize the protein^96,97^. Therefore it is likely that some of the translon-encoded peptides with non-canonical properties degrade quickly after their translation.

However, not all non-canoniocal peptides are necessarily short-lived. Stable non-canonical proteins have been identified whose knock-outs lead to specific phenotypic changes^5,22^. For example, peptides including Toddler/Elabella, and Pri are encoded by non-canonical translons and regulate embryonic development in zebrafish and Drosophila ^7,98–100^. Therefore, it is possible that some of the embryonic non-canonical translational events we identified may similarly affect mouse embryonic development. Identifying the molecular mechanisms and functions of these events will be a future direction.

Beyond their potential to encode functional micropeptides, upstream translons can also regulate gene expression by controlling ribosome loading on the CDS, ribosome stalling on the upstream translon, or activation of NMD^16,19–21,101^. We observed that ribosome occupancy of embryonic upstream translon was poorly but positively correlated with CDS occupancy in agreement with several previous studies^15,19,102^. Using RiboNN, a sequence-based deep learning model for predicting TE, we found that upstream translons predicted to reduce TE_CDS_ tend to initiate at ATG codons, whereas those predicted to increase TE_CDS_ initiate at near-cognate codons. To validate this, we mutated an ATG-initiating upstream translon in *Cnih2* that RiboNN predicted to be repressory, and confirmed reduced protein expression from an epitope-tagged exogenous construct.

We also predict that upstream translons in several ribosomal protein mRNAs might be regulatory. Rapid adaptation to environmental change via coordinated regulation of ribosomal protein mRNA expression and their translational buffering ^103,104^ is well known, and upstream translons might be an added layer that helps fine tune this regulation, especially if we can understand the role of these translons in specific stress conditions.

Finally, we highlight some of the important limitations of our study. First, while we observed subtle growth phenotypes upon disruption of select upstream translons, these effects are modest and suggest that many non-canonical translation events act in concert with other regulatory mechanisms rather than functioning as primary effectors. More fundamentally, because CRISPR-Cas9 screens generate a heterogeneous pool of indels, disentangling the contribution of translon perturbation from unintended coding sequence effects remains a significant interpretive challenge that will require allele-specific follow-up experiments. Second, although many of the identified translons are expressed during early development, establishing their roles in embryogenesis remains highly challenging, particularly because multiplex phenotypic assays in preimplantation embryos are technically demanding and the relevant developmental windows are narrow. Third, in this study, we did not pursue activity-dependent translational changes associated with LTP. Although our Ribo-ITP libraries from LTP-activated slices were technically reproducible, we observed highly variable expression patterns across timepoints, likely reflecting variability in the precise anatomical boundaries of microdissection. Comprehensive profiling of LTP-associated translation will therefore require more precise anatomical targeting and greater number of biological replicates.

Despite these limitations, we expect the utility of this approach to extend beyond the systems examined here. Recent studies have identified functional translons in diverse cancer cell lines^78,105–108^, and applying Ribo-ITP to rare tumor-resident cell populations could enable the discovery of novel tumor antigens as well as micropeptides that contribute to tumor maintenance and progression. Similarly, although many micropeptides have been detected through immunopeptidomic screens^4,109^, our approach offers a complementary strategy to identify translons expressed in antigen-presenting cells in response to infection. More broadly, integrating translon discovery from Ribo-seq with single-cell RNA-seq derived transcriptomic profiles across developmental stages may provide insights into cell-state and lineage-dependent regulation of non-canonical translation. Taken together, our results establish Ribo-ITP as a broadly applicable and scalable strategy for translon discovery, particularly suited to limited-input biological scenarios that have previously remained inaccessible.

## Materials and Methods

### Animals

All procedures were approved by the Institutional Animal Care and Use Committee at the University of Texas at Austin. C57BL6/J mice (Jackson Labs, RRID:IMSR_JAX:000664) were used for ribosome profiling experiments.

### Collection of mouse brain tissue for ribosome profiling

Mice were deeply anesthetized with a mix of ketamine (90 mg/kg; TW Medical, Decatur, GA) and xylazine (10 mg/kg; Animal Health International, Loveland, CO) intraperitoneally. Mice were transcardially perfused with ice-cold, oxygenated cutting solution, containing: 205mM sucrose, 25mM sodium bicarbonate, 2.5mM KCl, 1.25mM sodium phosphate, 7mM MgCl2, 7mM D-glucose, 3mM sodium pyruvate, 1.3mM ascorbic acid, and 0.5mM CaCl2. Brains were removed and hemisected, and the dorsal portion of the brain was removed using a scalpel blade, angled at 20°. The dorsal cut side was then mounted in the slicing chamber of a Leica VT1200 vibratome using superglue and submerged in ice-cold, oxygenated cutting solution. 250 mm oblique-horizontal hippocampal slices were made. Slices were incubated for 30 mins at 37°C in holding solution containing: 125mM NaCl, 25mM sodium bicarbonate, 2.5mM KCl, 1.25mM sodium phosphate, 12.5mM D-glucose, 2mM MgCl2, 2mM CaCl2, 1.3mM ascorbic acid, and 3mM sodium pyruvate. Slices were then stored for at least 30 mins in the holding solution at room temperature before being used for recordings. Reagents for electrophysiology solutions were purchased from Sigma Aldrich. Only slices from the middle of the dorsal-ventral axis of the hippocampus were used for recordings. Glass recording electrodes of 3–6 MOhms were pulled from thin-walled borosilicate filamented glass (Sutter BF150-110-10) using a P-1000 puller (Sutter). Electrodes were filled with artificial cerebrospinal fluid (ACSF) containing: 125mM NaCl, 25mM NaHCO3, 2.5mM KCl, 1.25mM NaH2PO4, 12.5mM D-glucose, 1mM MgCl2, and 2mM CaCl2. Data was acquired with a MultiClamp 700B amplifier (Molecular Devices) interfaced with a Digidata 1550B acquisition system and pClamp 11.1 software (Molecular Devices, San Jose, CA).

Slices were transferred to the recording chamber and continuously bathed in warm (32.5°C), oxygenated ACSF. Slices were visualized using a Zeiss Axio Examiner with Dodt contrast optics and epifluorescence, and a Zeiss Axiocam 503 infrared digital camera. Prior to recording, a small cut was made through the stratum radiatum at the border between CA3 and CA2 to reduce activation of recurrent connections. The recording electrode was placed in the stratum radiatum of CA1. Synaptic stimulation was delivered by a tungsten bipolar stimulating electrode (WE3ST30.1ASS, Microprobes) placed in the stratum radiatum near the CA2/CA1 border to stimulate the Schaffer collateral (SC) axons, and a constant current stimulus isolator (World Precision Instruments). First, slices were stimulated at 0.1 Hz, and field excitatory postsynaptic potentials (fEPSPs) were recorded for 10-30 mins to ensure that responses were stable, indicating good slice health. Next, the stimulus intensity was adjusted to evoke minimal and maximal responses and an input/output curve. The stimulus intensity was then set at the mid-point of this curve such that either increases or decreases in response could be recorded. Baseline fEPSPs were then stimulated and recorded at 0.1 Hz for 10 mins. Slices were then stimulated with a high frequency stimulus designed to induce LTP: 100 Hz for 1 s, repeated 4 times with a 20 s interval. Slices were then stimulated at 0.1 Hz for 10 mins, 30 mins or 60 mins, after which the slices were transferred to a new dish and the CA1 region was dissected under a dissection microscope. A slice that was approximately 1mm X 0.5mm was dissected using a sharp scalpel, transferred to a 1.5mL Eppendorf tube with minimal buffer carryover, flash-frozen in liquid nitrogen, and stored at −80°C.

The approximate number of neurons in the microdissected regions was estimated based on the volume of the microdissected region (1mm X 0.5mm X 0.2mm) and the neuronal density (80,000 cells/μm^3^) in CA1 region of mouse hippocampus ^110^.

### Mouse embryo isolation and treatments

Eight-week-old C57BL/6 female mice were injected with 5 U Pregnant Mare Gonadotropin (PMSG) (Ilex Life Sciences). Forty-eight hours later, they were injected with 5 U Human Chorionic Gonadotropin (HCG) (Millipore Sigma) and immediately paired with C57BL/6J male mice. Embryos were flushed from the oviducts on day 2.5 post-fertilization and cultured in KSOM medium (K0113, Cytospring). Approximately half of the embryos at the 16-cell stage were removed and remaining were cultured to the 32-cell stage. Embryos in both stages were separated equally into two experimental groups: KSOM + 2µg/mL Harringtonine and KSOM + DMSO (vehicle). The embryo groups were cultured for 30 mins, then rinsed through three drops of 20mg/mL BSA. Single embryos from each group were placed in thin-walled PCR tubes (VWR 20170-012) with minimal liquid volume using a pulled Pasteur embryo transfer pipette and flash-frozen in liquid nitrogen. The tubes were stored at −80°C.

### Ribo-ITP of microdissected mouse brain tissue and single embryos

Ribo-ITP was performed as described in Rao et al. ^111^. The brain tissue and single embryos were lysed differently as mentioned; the flash-frozen hippocampal tissue was thawed on ice, and 10μL of lysis buffer (1% Triton X-100, 50 mM MgCl2, 0.1 mg/mL cycloheximide) was added. The tissue was homogenized with a pestle until the solution was visibly clear. 4μL of the lysate was transferred to a 0.2mL PCR tube containing 1µL of 5X ITP lysis buffer (100mM BisTris pH 7.2, 5.0% Triton X-100, 25mM MgCl2, 12.5mM CaCl2, 500mM NaCl, 5mM DTT, 0.5mg/mL cycloheximide). For embryo lysis, flash-frozen single embryos were thawed on ice, 5µL of ITP lysis buffer (20mM BisTris pH 7.2, 1.0% Triton X-100, 5mM MgCl2, 2.5mM CaCl2, 100mM NaCl, 1mM DTT, 0.1mg/mL cycloheximide) was added to the droplet containing the embryo on the wall of the tube. For both the lysates, 1µL of RNaseI (Ambion) diluted 1:300 in the ITP lysis buffer was added and incubated at 37°C for 30 mins. 1µL of 0.7% SDS was added to stop RNaseI, followed by running the lysate on an ITP chip. Ribosome footprints were eluted by collecting the RNA fragments between 19nt and 36nt markers on the ITP chip. NGS libraries were made using Diagenode small RNA library preparation kit with the following modifications: a. 0.5µL PNK enzyme (NEB) was added along with the dephosphorylation reagent, b. The TSO was diluted 1:5, c. The cDNA was amplified for 16 cycles for embryos and 12 cycles for the hippocampal tissue. These cycle numbers were empirically tested and minimum cycle number yielding sufficient library was selected. The libraries were pooled and cleaned up with AmpureXP beads (1.5X) and purified on a 10% PAGE. The libraries were sequenced on a Novaseq 6000 (Illumina) instrument using the PE150 sequencing chemistry.

### Identification of translated ORFs

The ribosome profiling libraries were processed using the Riboflow pipeline^36^. The RiboR package was used for making metagene plots, region coverage plots and read length distribution and RiboGraph for data QC and visualization ^47,112^. For identification of translated ORFs from these transcriptome aligned files, all possible candidate ORFs were initially identified from the APPRIS release M25 (GRCm38.p6) transcriptome by extracting all start codons and pairing them with the first downstream in-frame stop codon. Only non-overlapping 3′UTR ORFs and non-overlapping and overlapping 5′UTR ORFs were considered. ORFs in the 5′UTR were limited to a length of 10 to 99 amino acids. For ORFs overlapping the CDS, only the portion within the 5′ UTR was considered for downstream analysis. RPFs were aligned and corrected for the ribosome P-site using an offset determined from the peak of the start site metagene profile within a range of 10-15 nucleotides. A dynamic cutoff algorithm [Citation error] was applied to include RPF lengths in which 85% of total CDS reads were contained. For the hippocampal dataset, ORF reads were aggregated across time points. For the embryo dataset, the 16-cell and 32-cell datasets were kept separate as two experiments. ORFs with fewer than 10 ORF reads per million experiment reads across all experiments, as well as those lacking significant 3-nucleotide periodicity based on chi-squared tests with false discovery rate (FDR) correction (*p* > 0.05) in all experiments, were excluded. To refine the set of in-frame ORFs within the same gene that share a stop codon, an additional filtering step was applied: longer ORFs were removed if all their reads were fully contained within a shorter in-frame ORF. Within a set of overlapping in-frame ORFs that share a common stop position, ORFs with non-canonical start codons were removed unless all ORFs in the overlapping group had non-canonical starts, in which case all were kept. The maximum CPM in each experiment was computed for each overlapping in-frame set of ORFs. Only ORFs containing at least 75% of this experiment-specific maximum in all experiments were retained, and if none met this threshold, all ORFs in the set were kept.

For genome alignment, the trimmed and mouse rRNA and tRNA filtered .fastq files obtained from Riboflow pipeline were aligned to the mouse genome (GRCm38/ mm10) using STAR (2.5.2) aligner with *end-to-end* (no soft-clip) and *--outSAMattributes All* mode. The aligned reads with a mapping quality cutoff of 10 were deduplicated using UMI tools version 1.1.1 with "--read-length" parameter. The genome-aligned sorted and indexed bam files were used to run RiboTISH 0.2.7^46^. First, the read distribution around the start and stop site, p-site offset, and the 3nt periodicity of the data was checked using *ribotish quality* with the regular riboseq option. These files were then used to identify all translons using the *ribotish predict* command with *--longest, --seq*, and *--aaseq* options. For the single embryo dataset, ‘*--altcodons CTG,GTG,TTG,ATT,ACG*’ and ‘--harr’ options were additionally used, while for the microdissected hippocampal dataset, only default options with *--longest* option was used.

### Amino acid composition comparison between translons and CDS regions

The amino acid composition between translons and CDS regions was compared with a Wilcoxon rank-sum test. The CDSs included were those of the same genes as the translons. The amino acid frequency was calculated for each translon and CDS, defined as the number of residues of each amino acid divided by the total number of residues. For each amino acid, a Wilcoxon rank-sum test was run, comparing the distribution of frequencies of the translon vs. CDS frequencies for the upstream and downstream translons. P-values were adjusted using Bonferroni correction. For visualization, the median frequency for each amino acid was calculated for the upstream and downstream translons, along with the CDSs. The percent change was calculated as the difference between the translon and CDS median divided by the CDS median for each amino acid.

### Translon expression across ribosome profiling studies

Uniformly processed .ribo files were obtained from RiboBase[Citation error], a curated repository of high-quality ribosome profiling datasets. All candidate ORFs were analyzed across available mouse experiments in RiboBase. For each Ribo-seq dataset, P-site offsets and read length cutoffs were dynamically determined as described above.

To ensure sufficient depth for detecting cell line–specific signals, datasets with fewer than 100,000 reads in the read length range were removed. To evaluate translation in specific cellular contexts, reads (CPM) were calculated for each ORF in each experiment and averaged across all experiments annotated to the same cell line.

To assess 3-nucleotide periodicity, datasets were further filtered by quantifying the number of CDS-mapped reads in each reading frame for each read length of an experiment [Citation error]. Experiments were removed if no read length had ≥75% of CDS-mapped reads aligned to a single reading frame. For each ORF, P-site coverage was aggregated across the remaining datasets. ORFs with a normalized read density below 0.1 reads per nucleotide were excluded. A one-dimensional discrete Fourier transform (DFT) was applied to the aggregated, P-site–adjusted coverage vector for each ORF. The frequency spectrum was computed using the *scipy.fft.fft* function, which calculates the DFT using the fast Fourier transform algorithm. The five frequency components with the highest amplitudes were identified, and their indices were normalized by the length of the frequency spectrum to yield relative frequencies ranging from 0 to 1. ORFs were retained if at least one of these five dominant frequency peaks fell within the 0.32–0.34 range, corresponding to 3-nucleotide periodicity.

This method determines 3-nucleotide periodicity by combining two complementary strategies: filtering based on global frame mapping along with a frequency-domain analysis of periodicity across the full read coverage profile.

### Correlation of ribosome occupancy of translons vs. CDS

To evaluate the relationship between translon and main CDS ribosome occupancy, we analyzed CPM-normalized paired Ribo-seq and RNA-seq read counts from RiboBase. A partial Spearman correlation was performed using the R *ppcor* package to compare translon CPM and CDS CPM values, while controlling for gene-level RNA-seq expression to account for differences in transcript abundance. Correlations were calculated separately for hippocampal and embryonic datasets, and for upstream and downstream translons within each. To assess whether the resulting correlation coefficients (Spearman’s ρ) were significantly different from zero, we performed a Wilcoxon signed-rank test on the ρ values in each group.

The CPM values of each translon averaged across all datasets within each cell line were used to compute a Gini coefficient which determines the cell-line specificity of each translon. The Gini coefficient was calculated with the R *DescTools* package^113^. to assess whether cell-line specificity was associated with translon-CDS translation relationship, Spearman correlation was calculated between the Gini coefficients and the partial Spearman correlation coefficients for upstream and downstream translons. Translons containing no RPF reads in the CDS region in all datasets were excluded.

### Cell culture

HEK293T cells were cultured in Dulbecco’s Modified Eagle Medium (DMEM) with 10% fetal bovine serum (FBS) and 1% penicillin-streptomycin. C57BL/6 mouse embryonic stem cells were obtained from ATCC, acclimatized to non-feeder culture conditions, and later grown in 2i medium (Knockout DMEM with 15% FBS, 100 U/mL penicillin-streptomycin, 2mM glutamine, 1X non-essential amino acids, 0.15mM 2-mercaptoethanol, 100U/mL Lif, 3 μM CHIR99021, 1μM PD0325901)

### Cloning and expressing of GFP and Cnih2 reporter constructs

The GFP sequence with (startGFP) and without (nostartGFP) the start codon was amplified and inserted into the pLVX vector (Addgene #141367) between the BamHI and MluI restriction sites using In-Fusion® Snap Assembly (Takara). In the nostartGFP plasmid, all 5′UTRs were cloned in-frame with the nostartGFP (between the EcoRI and BamHI sites) using In-Fusion® Snap Assembly. Insulin 5′UTR along with the CDS was cloned as a positive control. The 5′ UTR sequences of the seven transcripts harbouring translons were cloned until the stop codon of the translon to test their translation capacity. Actin 5′UTR, which did not have any predicted translon, was cloned in both StartGFP and nostartGFP plasmids. Sequence-confirmed individual plasmids were reverse transfected in a single cell suspension of mESCs with Lipofectamine 2000 (Invitrogen). Live cells were imaged using a STELLARIS 8™ fluorescence microscope (Leica) 24 hours post-transfection. The brightness and contrast of the images were adjusted post-imaging for effective visualization.

The Cnih2 transcript was cloned in pLVX_mpuro (Addgene #141367) vector between XhoI and EcoRI sites using In-Fusion® Snap Assembly (Takara). Cnih2 transcript was amplified as three fragments, first from start of the 5′UTR until the stop codon of the upstream translon with an inframe Flag tag, second from the stop codon of the upstream translon until the stop codon of Cnih2 with an inframe HA tag and third from stop codon of Cnih2 until the end of the 3′UTR. This plasmid was used as template for SDM of the upstream translon ATG to AGG. Both the WT (ATG) and mutant (AGG) plasmids were sequence confirmed and transfected in HEK293T cells with Lipofectamine 3000 using the manufacturer’s protocol and used for either western blot analysis or imaging analysis. For western blot analysis, the transfected cells were washed with PBS and lysed in 200μL RIPA buffer with protease inhibitor cocktail and centrifuged at 10,000g for 10 mins. Protein concentration of the clarified lysates was confirmed with Qubit Protein Broad Range assay kit. Equal amounts of lysate were loaded on a Bolt 4-12% Bis-Tris protein gel and transferred onto a 0.2μ nitrocellulose membrane. Membranes were blocked in 5% milk and incubated with mouse anti-FLAG (Sigma-F3165, 1:4000), rabbit anti-HA (Abcam ab9110, 1:4000), and mouse anti-Actin (MP Bio- 0869100-CF, 1:500) primary antibodies and Goat Anti-Rabbit IgG H&L (HRP) (ab6721) and Goat Anti-Mouse IgG H&L (HRP) (ab205719) secondary antibodies. Blots were developed with SuperSignal Pico Plus substrate, visualized with Biorad ChemiDoc MP and analyzed on the BioRad ImageLab software and the data plotted using R. For imaging analysis, transfected cells were washed thrice with 1x PBS, fixed with 4% PFA dissolved in 1x PBS for 10mins, washed with 1x PBS twice and blocked in 5%FBS in 0.1% PBS-T for 1 hour at room temperature. Cells were incubated with the mouse anti-FLAG and rabbit anti-HA antibodies mentioned above for 1 hour at room temperature followed with 3 washes with 0.1% PBS-T and incubation with Goat anti-Mouse IgG (H+L) Alexa Fluor Plus 647 (Invitrogen A32728) and Donkey anti-Rabbit IgG (H+L) Alexa Fluor Plus 488 (Invitrogen 32790), washed thrice with 0.1% PBS-T. 1µg/Ll DAPI was added to the final wash and slides were mounted in VECTASHIELD® Vibrance™ Antifade Mounting Medium, With DAPI, Liquid (H-1800-2). Slides were imaged using a STELLARIS 8™ confocal fluorescence microscope (Leica). .lif files were analyzed in Fiji by manually drawing ROIs for individual cells using the DIC image and quantitating the fluorescence for the ROIs in the 488 channel. Fluorescence values were plotted using R. The brightness of the images were adjusted post-imaging for effective visualization.

### Large-scale GFP reporter screen

To perform a large-scale GFP reporter screen all translons (hippocampal + embryonic) that were less than 250bp in length were selected. Then 250bp 5′UTR sequence upstream of the translon stop codon was selected from the Appris transcript fasta sequence. If the final sequence was less than 100bp in length, it was filtered out and all the remaining sequences (100bp-250bp) were synthesized as a pool and cloned in the pLVX-nostartGFP vector by Twist Biosciences as previously described. To enable puromycin selection, this library containing the 5′UTR sequence in-frame with the nostart-GFP sequence was further subcloned in a lentiviral pBOB-EF1-PGK-Puro vector (Addgene #86849) between NotI and XhoI enzyme sites. This pooled guide library was then used for generating lentivirus. HEK293T cells were grown in T75 flasks and transfected with equimolar ratio of psPAX2, pMD2.G, and the pBOB-translon pooled library plasmids using Lipofectamine 3000 reagent (Invitrogen) according to manufacturer’s protocol. Viral titer was calculated by infecting mES cells with serial dilutions of virus followed by puromycin selection and colony counting.

For screening, 2.5 x 10^6^ mESCs were infected with the lentiviral library at an MOI of 0.3 and a library representation of 500X in T75 flask overnight. Next day, the media was replaced with 2i ES growth media, and cells were allowed to grow for 24 hours. Infected cells were selected using 2i ES growth media + 0.3µg/mL puromycin for the next 48 hours. The surviving cells were recovered in 2i ES growth media for next 48 hours. Following recovery cells were trypsinized and resuspended in 1mL PBS+ 1% FBS to reduce clumping. 100µL of the cells were stored as input and remaining 900µL cells were passed through a cell strainer to achieve single cell suspension and immediately sorted using a Sony MA900 FACS machine using the maximum purity settings. Singlets were gated using a SSC-H vs SSC-W plot followed by sequential gating using a FSC-A vs FSC-H plot. Singlet cells showing a 10 fold increase in GFP were considered GFP positive and sorted in PBS+ 5% BSA to increase centrifugation efficiency. Sorted cells were centrifuged and the pellets were used for genomic DNA isolation using Zymo mini DNA isolation kit. The 5′UTR sequences upstream of the nostartGFP were amplified from 2 µg gDNA in a 100µL PCR reaction divided into four 25 µL reactions for 22 cycles using custom made Illumina indexing primers to generate amplicon libraries (Supplemental table S8). PCR products were cleaned with AmpureXP beads and quantified on a bioanalyzer. Guide amplicon libraries were sequenced using NovaSeq 6000 (Illumina) with a PE150 chemistry.

### Analysis of Large scale GFP reporter screen

NGS reads were trimmed to remove vector-specific PCR handles using Cutadapt^114^ v1.18 with the following parameters: “-g ^CGTGAGGATCTATTTCCGGTGAATTC -G ^GGTGAACAGCTCCTCGCCCTTGCTGGATCC”. Trimmed reads were aligned to the APPRIS release M25 (GRCm38.p6) transcriptome using bowtie2^115^ v2.3.4.3. The resulting sorted indexed bam files were used to generate a count table using a custom script. The raw counts for the Input and GFP+ samples were CPM normalized and mean log2 expression (M) and mean log2FC (A) were calculated across replicates. Translons which had consistent mean log2FC > 2 and mean log2 expression > 2 in both the replicates were considered as enriched in GFP+ population compared to the input. Different translon features (type, start codon preference, length) were compared between the enriched vs. non-enriched translons. All the plots were made using the ggplot2 package in R v2024.04.2+764.

### RiboNN predictions and analysis

The chi-square filter passed hippocampal and embryonic translons were used for this analysis. A. The RiboNN pre-trained mouse model was installed and ran according to the directions in the provided GitHub repo ^83^. For each transcript with a translon of interest, multiple mRNA sequences were generated: a wild type sequence, a sequence with the start codon of the translon mutated to a stop codon, and five control sequences where random positions in the 5′UTR or 3′UTR (for upstream and downstream translons respectively) that do not fall into any other known ORFs were mutated to a stop codon. This process was repeated for each of the three stop codons and compiled into the input format required by RiboNN. Predicted TE was calculated as the mean over all mouse cell types predicted by the model.

The .csv file containing the predicted TE values (Supplemental table S9) for all the translons across the different conditions was transferred to R v2024.04.2+764. delTE was calculated by subtracting TE_mut_ from TE_wt_. A mean delTE value was calculated for the five controls and the three stop codons. Wilcoxon’s sum rank test was used for significance testing between distribution of meandelTE values of the Start mutation and control. The top 20 high delTE and low delTE values were selected, translon type and start codon composition were identified. All plots were made using the ggplot2 package.

### Translon Pooled Knockout Screen

The genomic locations of the identified translons were used to design guide RNAs against the targets using CHOPCHOP v3 ^116^ and IDT’s CRISPR-Cas9 guide RNA design tool. The following criteria were used for choosing the best 2 guides per translon: high on-target score, low mismatch score, and position on the predicted upstream translon In case of overlapping ORFs, we chose guides which cut as far as possible from the annotated ORF. The list of guide sequences used is mentioned in Supplemental table S8. These guides were cloned into lentiCRISPR v2 (Addgene #52961) as a pool, as described for the GeCKO Lentiviral CRISPR Toolbox ^117^. A list of the guides and oligos cloned for this screen is provided in Supplemental Table S8. Briefly, first the lentiCRISPRv2 vector plasmid (Addgene # 52961) was digested with BsmBI-v2 (NEB) for 1 hour at 37°C and heat-inactivated at 80°C for 10 mins. The digested product was separated on a TBE-Agarose gel, and a 8kb band was cut and purified using a gel extraction kit (Macherey-Nagel). The forward and reverse oligos for each guide sequence with flanking BsmBI sites (IDT) were phosphorylated with T4 PNK (NEB) and hybridized as individual pairs by incubating at 37°C for 30 mins and inactivation at 95°C for 10 mins, followed by ramping down the temperature to 4°C at a rate of 5°C/min. Equimolar quantities of all the hybridized guides were pooled together and ligated with 1 μg linearised lentiCRISPR v2 plasmid at room temperature for 30 mins. Ligated product was transformed in 250 µL of chemically competent NEB Stbl3 cells by heat shock at 40°C for 1 min followed by incubation on ice for 2 mins and recovery in 10 mL warmed SOC outgrowth medium (NEB) for 1 hour. All cells were plated on a bioassay plate and incubated at 37°C overnight. Transformants were collected by scraping all the colonies off the plate in LB+Amp, and plasmid was isolated using a plasmid maxiprep kit (Zymo) including endotoxin removal steps. This pooled guide library was then used for generating lentivirus. HEK293T cells were grown in T175 flasks and transfected with equimolar ratio of psPAX2, pMD2.G, and the pLentiCRISPRv2_pooled library plasmids using Lipofectamine 3000 reagent (Invitrogen) according to manufacturer’s protocol. Viral titer was calculated by infecting mES cells with serial dilutions of virus followed by puromycin selection and colony counting.

1.3 x 10^5^ mESCs were infected in a T25 flask with the pooled sgRNA lentivirus library at a multiplicity of infection of 0.3 such that there was >100X representation of each guide. Next day, the media was replaced with 2i ES growth media, and cells were allowed to grow for 24 hours. Infected cells were selected using 2i ES growth media + 0.3µg/mL puromycin for the next 48 hours. The surviving cells were seeded in a T75 flask (considered as Day 1 for the screen) and kept on selection for 8 days followed by recovery in 2i medium for 2 days. A portion of Day 1 cells was stored at −20°C as input. After 10 days of growth (approximately 20 cell division cycles), cells were harvested and genomic DNA was isolated using Zymo mini DNA isolation kit. Genomic regions with the guide sequences were amplified from 5 µg gDNA in a 100µL PCR reaction divided into four 25 µL reactions for 22 cycles using custom made Illumina indexing primers to generate amplicon libraries (Supplemental table S8). PCR products were cleaned with AmpureXP beads and quantified on a bioanalyzer. Guide amplicon libraries were sequenced using NovaSeq 6000 (Illumina) with a PE150 chemistry.

### Analysis of upstream translon pooled knockout screen

NGS reads were trimmed to remove vector-specific PCR handles using Cutadapt^114^ v1.18 with the following parameters: “-g ACACTCTTTCCCTACACGACGCTCTTCCGATCTGTGGAAAGGACGAAACACCG -a GTGACTGGAGTTCAGACGTGTGCTCTTCCGATCTCCAATTCCCACTCCTTTCAAGACC”. Trimmed reads were aligned to all guide RNA sequences used in the screen using bowtie2^115^ v2.3.4.3. A count table was made using a custom script which was used to calculate a ratio of guide abundance on day 1 to guide abundance on day 10. Further, MAGeCK analysis was also performed to identify significantly under and over represented guides using the count table with the following parameters: --paired --adjust-method fdr and --sort-criteria negative.

## Data Availability

All ribosome profiling raw data sets and the processed .ribo files generated in this study have been submitted to the Gene Expression Omnibus GSE300869, GSE300832, GSE318608. All custom scripts used to perform bioinformatics analyses are available at GitHub (https://github.com/up114/translons) and as Supplemental Code.

## Acknowledgements

We thank W. Shawlot and the UT Austin Mouse Genetic Engineering Facility for their contributions. This work was supported by the National Institutes of Health grants (R35GM150667; HD110096), as well as a Welch Foundation grant (F-2027-20230405) (C.C.). Figures were generated using BioRender.com under a publication license (JO26GB978U). All original text in this paper was authored by the researchers. Additionally, we acknowledge the assistance of a Large Language Model (OpenAI) for suggesting edits aimed at improving clarity and grammar.

## Author contributions

V.G. wrote the original manuscript. U.P., M.H., and C.C. participated in reviewing and editing the manuscript. V.G. and C.C. conceptualized the study design. V.G. and U.P. generated the figures for the manuscript. V.G. performed all the ribosome profiling experiments, quality control analysis of the libraries, GFP screen, Crispr screen and delTE analysis and validation. L.P. performed the RiboNN predictions. Y.T. performed cloning and western blots. M.H. performed electrophysiology recordings, U.P. developed the transcriptomic analysis pipeline and RiboBase analysis workflows. C.C. provided study oversight and acquired funding. All authors approved the final manuscript.

