## Supplemental figures for "Ribo-ITP enables identification of translons from limited input samples"

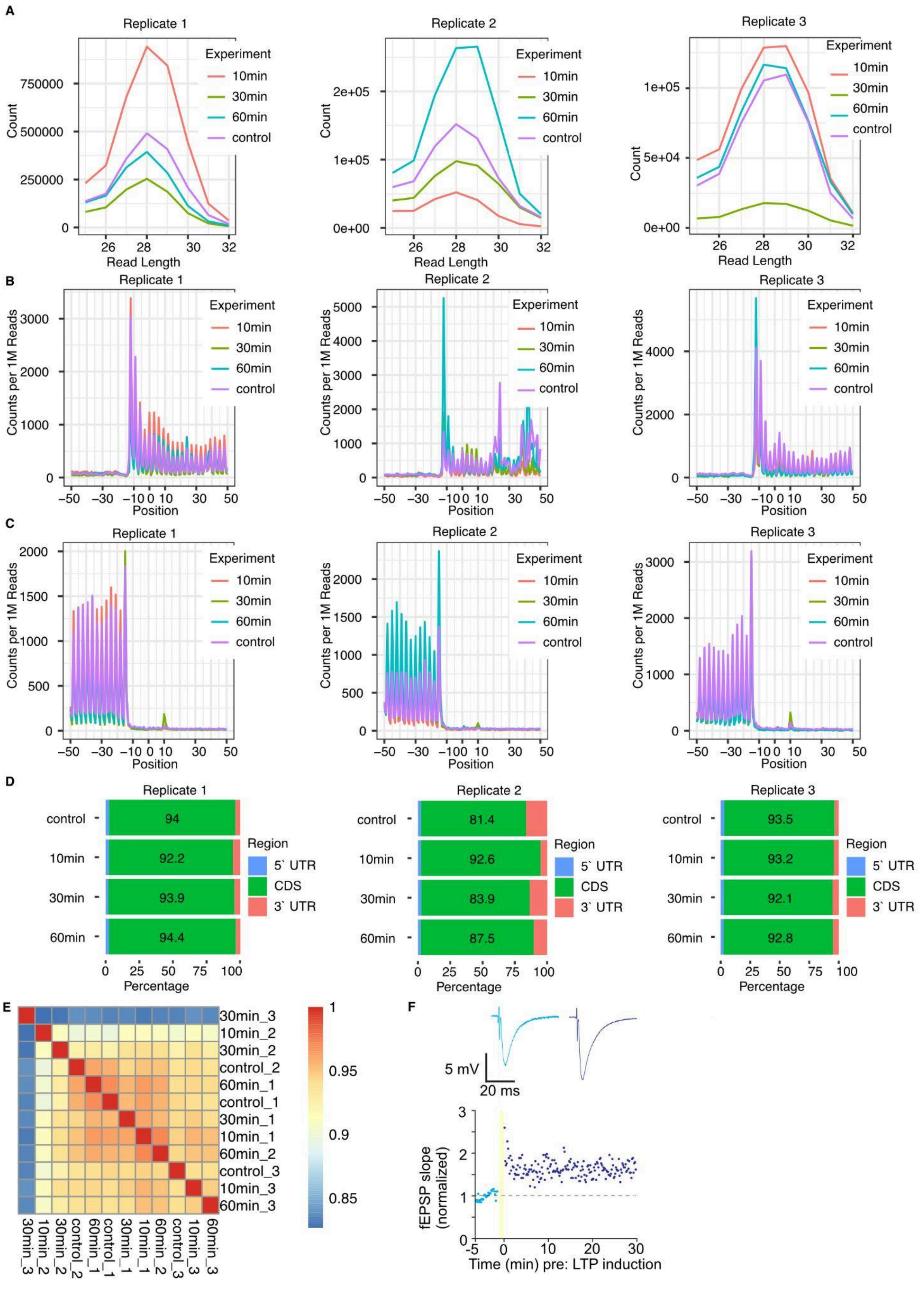

FigS1: QC plots for the individual hippocampal Ribo-ITP replicates:

*A, Read length distribution plots of the transcriptome mapping RPFs from individual biological replicates. The raw read counts for each read length are plotted. B, The Ribosome occupancy around the translation start site for individual replicates. Translation start site is denoted by the position 0. Aggregated read counts (y-axis) relative to the start site are plotted without P-site offset correction. C, The Ribosome occupancy around the translation stop site for individual replicates. Translation stop site is denoted by the position 0. Aggregated read counts (y-axis) relative to the stop site are plotted without P-site offset correction. D, Read coverage across different regions of the transcript. Percent mapping reads in the 5'UTR, CDS and 3'UTR are plotted. E, Replicate similarity between the four time points across the three biological replicates. Heatmap of the Spearman's correlation R values for all the hippocampal datasets. F, Voltage traces of LTP induction in hippocampal slices. Plots on the top show traces recorded before (light blue) and after (dark blue) the presentation of the LTP stimulus. The plot on the bottom shows the slope of the response as a function of time. The yellow shaded region is the time during which the LTP stimulus is being presented.*

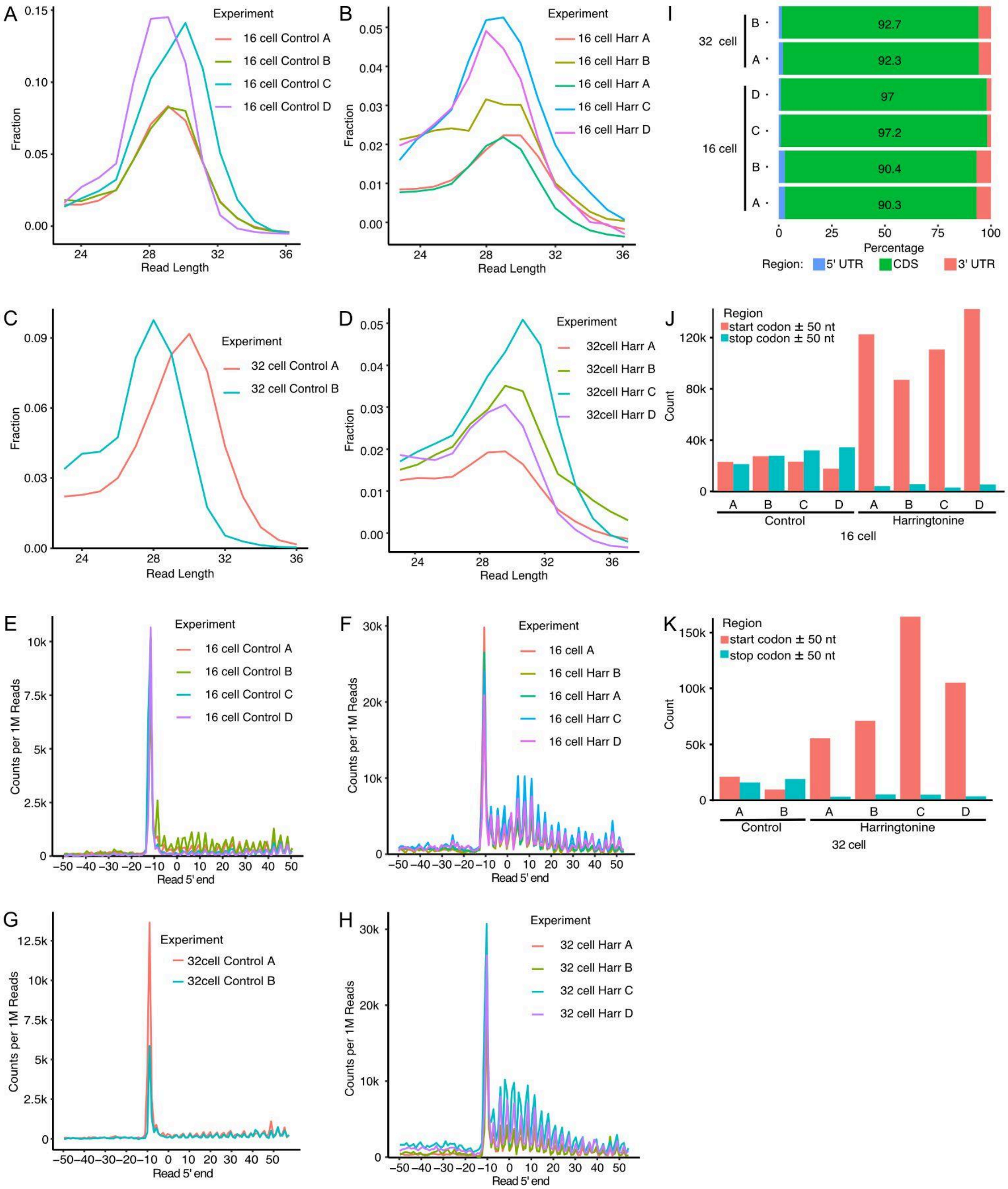


FigS2: QC plots for the individual embryo Ribo-ITP replicates:

*A-D, Read length distribution plots of the transcriptome mapping RPFs from individual embryos. Normalized read counts for each read length are plotted. E-H, Ribosome occupancy around the translation start site for individual embryos. Translation start site is denoted by the position 0. Aggregated read counts (y-axis) relative to the start site are plotted without P-site offset correction. I, Read distribution plot showing the percent reads mapping to the different regions in a transcript (5'UTR, CDS and 3'UTR). J-K, Distribution of raw reads mapping in the 50 nt region surrounding the translation start site and stop site for individual embryos.*

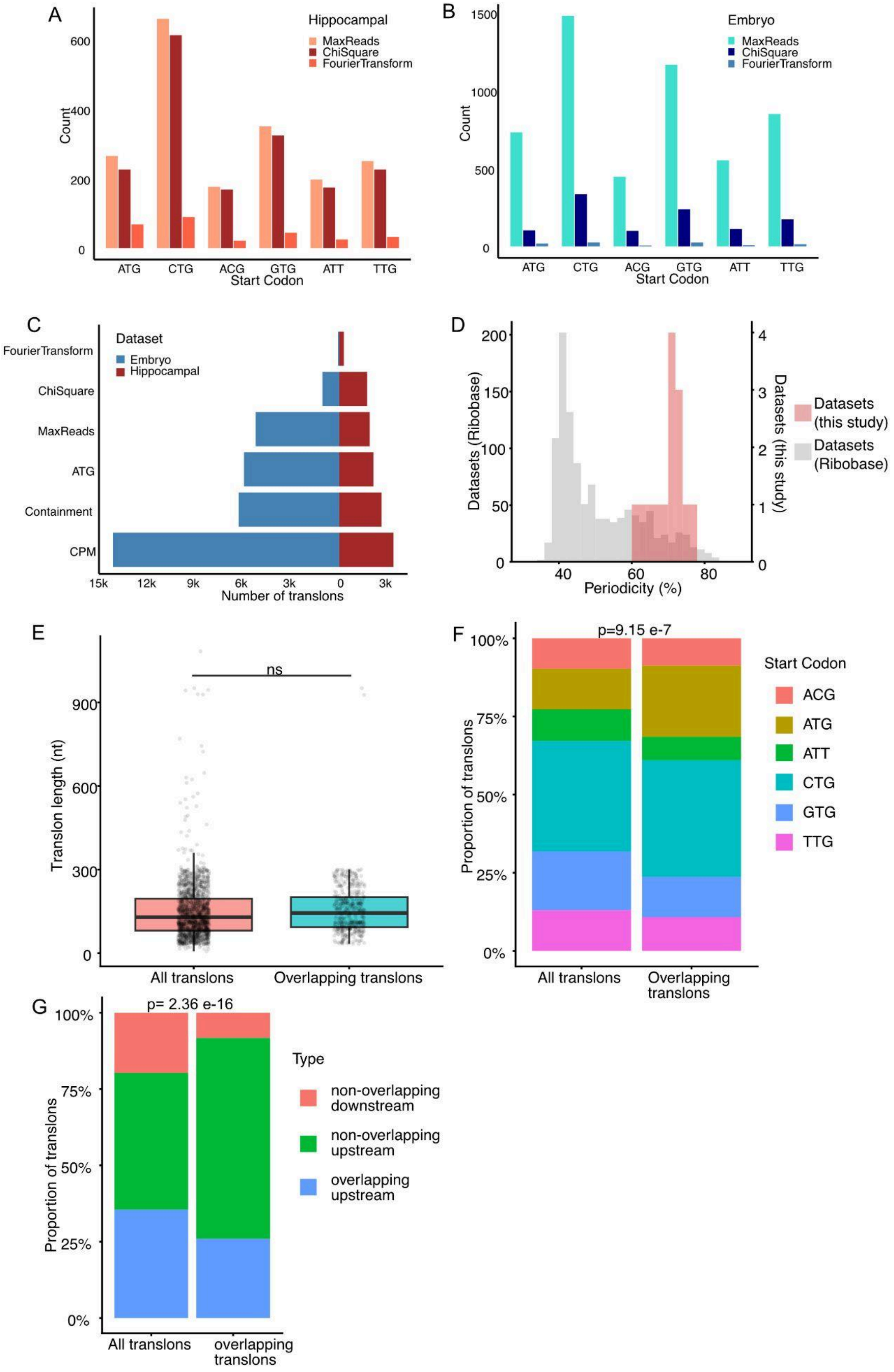

A
Hippocampal
MaxReads
ChiSquare
FourierTransform
Count
0
200
400
600
ATG
CTG
ACG
GTG
ATT
TTG
Start Codon
B
Embryo
MaxReads
ChiSquare
FourierTransform
Count
0
500
1000
1500
ATG
CTG
ACG
GTG
ATT
TTG
Start Codon
C
Dataset
Embryo
Hippocampal
FourierTransform
ChiSquare
MaxReads
ATG
Containment
CPM
15k
12k
9k
6k
3k
0
3k
Number of translons
D
Datasets (Ribobase)
Datasets (this study)
Datasets (this study)
Datasets (Ribobase)
0
50
100
150
200
0
1
2
3
4
40
60
80
Periodicity (%)
E
ns
Translon length (nt)
0
300
600
900
All translons
Overlapping translons
F
p=9.15 e-7
Proportion of translons
0%
25%
50%
75%
100%
Start Codon
ACG
ATG
ATT
CTG
GTG
TTG
All translons
Overlapping translons
G
p= 2.36 e-16
Proportion of translons
0%
25%
50%
75%
100%
Type
non-overlapping downstream
non-overlapping upstream
overlapping upstream
All translons
overlapping translons

*FigS3: Translon quality control*

*A, Histogram showing the start codons of hippocampal translons that pass the three different filters in the transcriptomic pipeline. B, Histogram showing the start codons of embryonic translons that pass the three different filters in the transcriptomic pipeline. C, Bar plots showing the number of translons that pass the three different filters in the transcriptomic pipeline. Colored in blue are embryonic translons and colored in maroon are the hippocampal translons. D, Histogram showing the periodicity distribution of studies in ribobase in grey and ribo-seq libraries generated in this study in salmon. E, Box plots showing the length of all the translons compared to common translons identified by both genomic and transcriptomic pipelines. Significance was tested using Wilcoxon's sum rank test. F, Plots showing the proportion of start codons of all the translons and common translons identified by both genomic and transcriptomic pipelines. G, Plots showing the proportion of translon types in all the translons and common translons identified by both genomic and transcriptomic pipelines.*

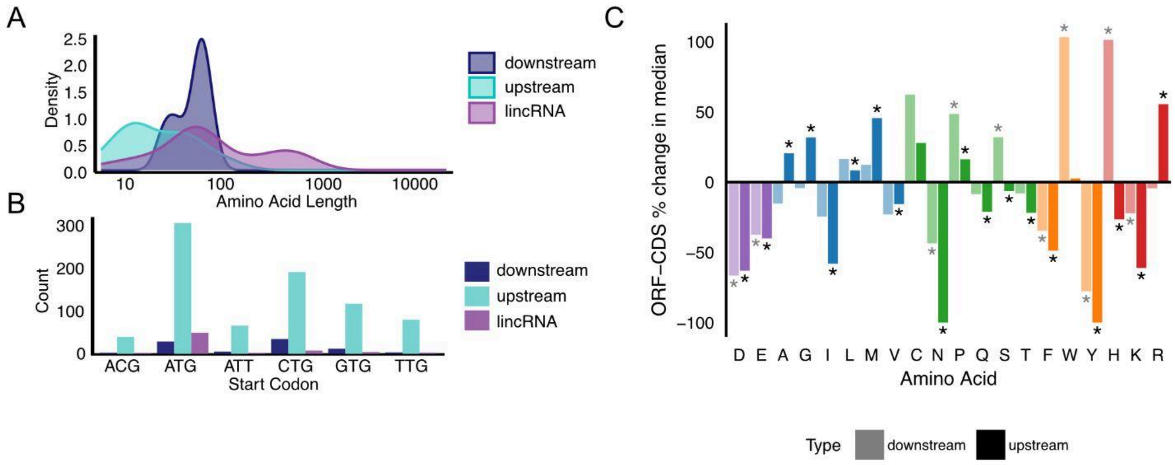


*FigS4: Properties of translated non-canonical ORFs identified by RiboTISH*

*A, Density plots showing the amino acid length distribution of the peptides encoded by the translons identified in the hippocampal dataset identified by RiboTISH. B, The number of translons identified with cognate and near-cognate start codons in the hippocampal dataset identified by RiboTISH. C, Bar graph showing the fractional difference in the upstream (dark) or downstream (light) hippocampal translons identified by RiboTISH with their corresponding CDS for the individual amino acids. Colors indicate amino acid type. * indicates Wilcoxon's rank-sum test p values < 0.05.*

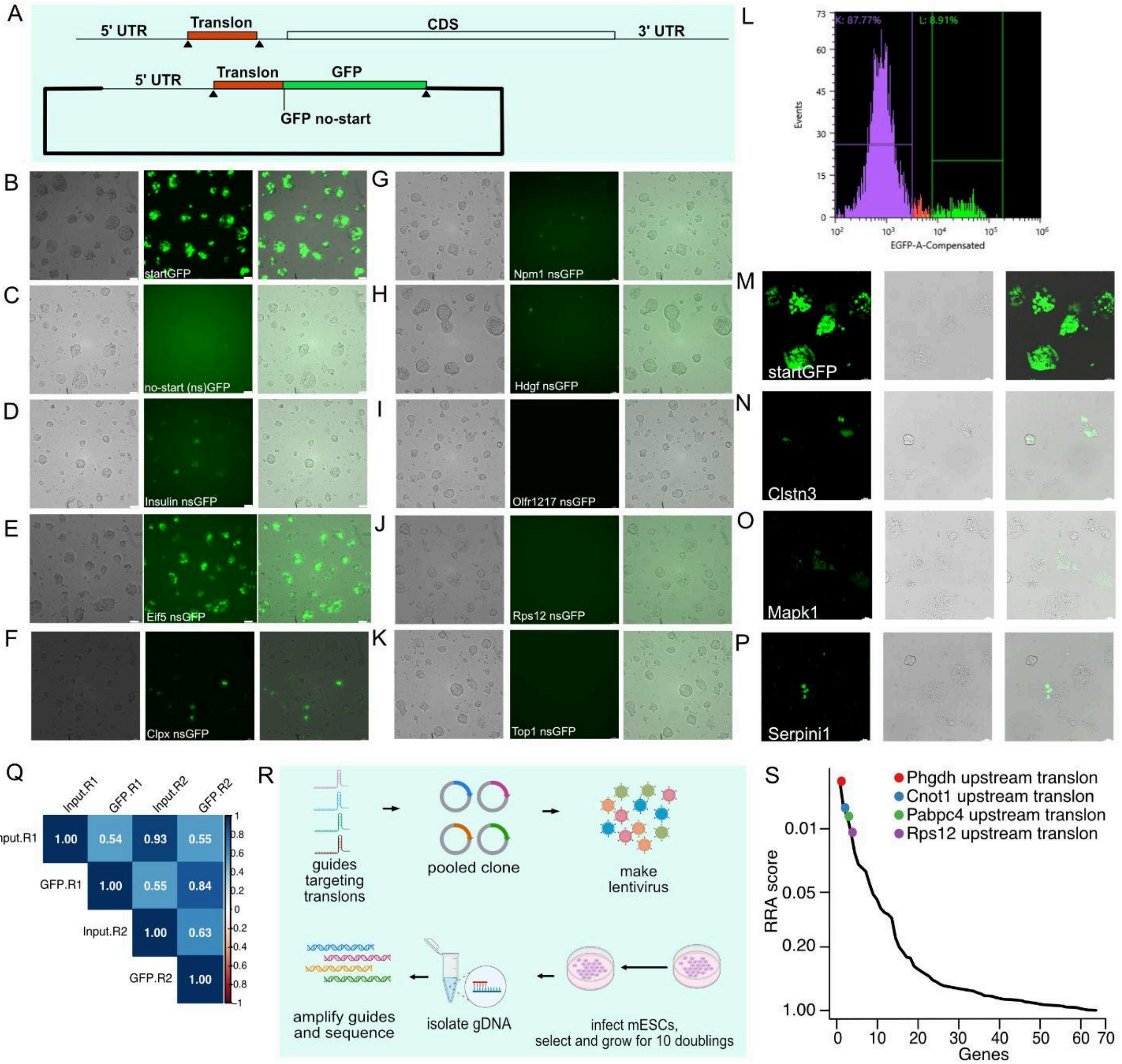

*FigS5: Non-canonical translons can drive translation of a reporter and affect cellular proliferation A, Schematic of the reporter design for GFP based translational capacity reporter. B-K, Fluorescence, DIC and merged images of mESCs transfected with the 5'UTRs containing the candidate ORFs cloned in the GFP reporter. Scale bar width is adjusted for effective visualization purpose, scale bar = 50µm. L, Representative histogram showing GFP+ and GFP- cell populations during FACS sorting for the high throughput screen. The population shown in Green was gated GFP+ and sorted. M-P, Fluorescence, DIC and merged images of mESCs transfected with clones tested in the high-throughput screen. Q, Spearman correlation pot showing the pairwise correlation between the high throughput GFP screen replicates. R, Schematic of the CRISPR screen experimental setup. R, Histogram of the RRA scores for upstream translons tested. The highlighted ORFs have a low RRA score with a FDR less than 25%.*

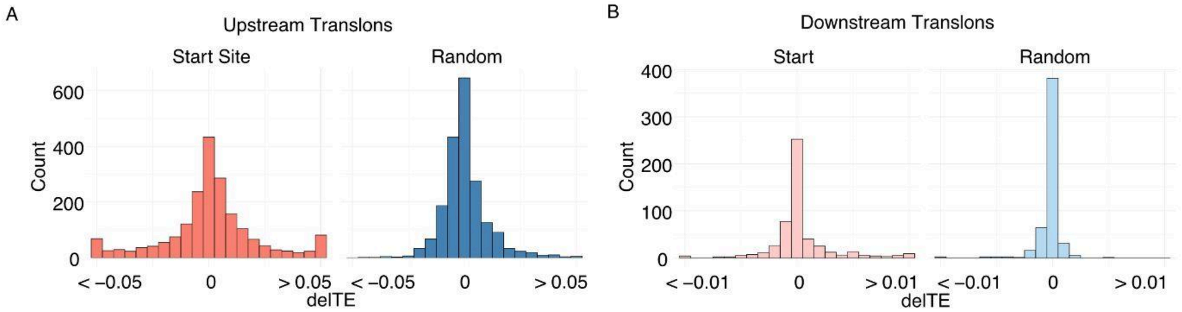


*FigS6: Effect of translon start codon mutation on CDS expression*

*A, Histograms of the predicted delTE values of the CDS upon mutating the start site of the translon or a random site in the 5'UTR outside the translon to a stop codon. Values at beyond the x-axis limits are added to the last quantile. B, Histograms of the predicted delTE values of the CDS upon mutating the start site of the translon or a random site in the 3'UTR outside the translon to a stop codon. Values at beyond the x-axis limits are added to the last quantile.*
